# ShuTu: Open-Source Software for Efficient and Accurate Reconstruction of Dendritic Morphology

**DOI:** 10.1101/226548

**Authors:** Dezhe Z. Jin, Ting Zhao, David L. Hunt, Rachel P. Tillage, Ching-Lung Hsu, Nelson Spruston

## Abstract

Neurons perform computations by integrating inputs from thousands of synapses – mostly in the dendritic tree – to drive action potential firing in the axon. One fruitful approach to understanding this process is to record from neurons using patch-clamp electrodes, fill the recorded neuron with a substance that allows subsequent staining, reconstruct the three-dimensional architecture of the dendrites, and use the resulting functional and structural data to develop computer models of dendritic integration. Accurately producing quantitative reconstructions of dendrites is typically a tedious process taking many hours of manual inspection and measurement. Here we present ShuTu, a new software package that facilitates accurate and efficient reconstruction of dendrites imaged using bright-field microscopy. The program operates in two steps: (1) automated identification of dendritic process, and (2) manual correction of errors in the automated reconstruction. This approach allows neurons with complex dendritic morphologies to be reconstructed rapidly and efficiently, thus facilitating the use of computer models to study dendritic structure-function relationships and the computations performed by single neurons.

**Significance Statement:** We developed a software package – ShuTu – that integrates automated reconstruction of stained neurons with manual error correction. This package facilitates rapid reconstruction of the three-dimensional geometry of neuronal dendritic trees, often needed for computational simulations of the functional properties of these structures.

## Introduction

The geometry of dendritic arbors directly influences synaptic integration and the resultant firing patterns of neurons (Mainen and Sejnowski, 1996; Henze et al., 1996; Stuart and Spruston, 1998; Krichmar et al., 2002). Dendritic morphologies vary widely across and within regions of the brain (Parekh and Ascoli, 2013), so consideration of morphology is an important aspect of understanding the mechanisms by which different neurons carry out their unique functions. Intracellular recording of neurons is a common technique for studying dendritic integration of input signals (Hamill et al., 1981; Stuart and Spruston, 1998). To fully understand the implications of these experiments, numerical simulations of the recorded neurons are often needed (Jaeger, 2001; Krichmar et al., 2002; Gidon and Segev, 2012; Menon et al., 2013). Informative simulations require accurate reconstructions of the geometry of the recorded neurons, including branching structures and diameters of the branches.

The traditional method of reconstructing neuron morphology requires intensive human labor (Zandt et al., 2017). A slide containing a neuron filled with biocytin is mounted on a motorized stage and imaged using a video camera mounted to a bright-field microscope. The neuron image is displayed on a computer screen, and the reconstruction is done manually. The user clicks the mouse along the images of dendritic branches on the screen. While clicking, the user adjusts the cursor size to match the diameters, and turns the z-position knob on the microscope to keep the branches in focus. Each click records the *x*, *y*, and *z* positions and the radius *r* at a single point, and connects the point to the previously clicked point. Bifurcations are marked and followed up sequentially. The morphology is recorded in a series of these clicked points.

Manual reconstruction in this way is computationally straightforward. Since it requires no image storage or processing, the computational demand is minimal. However, there are several drawbacks, especially when the accuracy of reconstruction is crucial. Repetitive clicking while measuring the radii and turning the focus knob makes manual reconstruction labor-intensive and time-consuming. The problem is exacerbated at high magnification. To see fine processes of neurons, it is desirable to image neurons with an objective at 100X magnification and a large numerical aperture (Jaeger, 2001; Brown et al., 2011). In our experience, however, it can take 10 − 15 hours or more of continuous work to reconstruct the dendritic tree of a pyramidal neuron in this way. Over this period of time, instability of the sample in the microscope can lead to problems. Furthermore, the accuracy of the reconstruction can suffer from fatigue-induced mistakes. Another problem with manual reconstruction is that the accuracy is hard to check independently because it is difficult to precisely align the previous reconstruction with the neuron image after remounting the slide.

Automatic reconstruction of neuron morphology using computer algorithms promises to reduce manual labor and increase productivity. There have been intensive efforts towards this goal, including open-source projects such as the Digital Reconstruction of Axonal and Dendritic Morphology Challenge (DIADEM)(Liu, 2011; Svoboda, 2011; Gillette et al., 2011; Gillette et al., 2011) and the BigNeuron project (Peng et al., 2015). Commercial software is also moving in this direction. In our experience, however, available software suffered from a variety of problems, including limited automation and tedious approaches for error correction. Thus, we sought to develop an open-source software platform that would overcome these limitations. In this paper, we describe our open-source software package, ShuTu (Chinese for “dendrite”) – a system for reconstructing biocytin-filled neurons efficiently and accurately. To avoid the impression of marketing our software, we make no attempt to compare it to other open-source or commercial software; instead, we encourage others to try it and judge for themselves.

## Results

We demonstrate the use of ShuTu by going through the steps involved in reconstructing a single CA3 pyramidal neuron from a mouse hippocampal slice. We then present reconstruction results for other cell types as well. Neurons were stained following patch-clamp recordings in brain slices prepared from 17-30 day-old male mice (C57Bl/6), using biocytin-containing intracellular solution. Following recording and staining, neuron reconstruction proceeded according to the following steps: (1) image acquisition; (2) image processing; (3) automated reconstruction; (4) manual editing and error correction. Additional details regarding slice recordings and computer systems requirements are provided in the Materials and Methods section. Operational commands for ShuTu are provided in Appendix 1. Technical details regarding the algorithms used in ShuTu are provided in Appendix 2.

### Image acquisition

ShuTu uses tiles of tiff stacks covering the entire neuron (Fig. 1). Nearby tiles should overlap by *∼*20%, in order to facilitate accurate stitching of tiles into a single image. We imaged hippocampal neurons using a Zeiss AxioImager microscope with AxioCam and ZEN blue software. Once the boundary in the field of view (*xy*) and the range of the depths (*z*) that contain the neuron were set, the images at each tile position and depth were acquired automatically, and the positions of the these images were stored in an xml file (image metadata). Other microscope/software combinations can be used, as long as tiff stacks and their relative positions are provided to ShuTu (see Materials and Methods for details). It is also possible to use ShuTu to reconstruct neurons imaged using two-photon, confocal, or wide-field fluorescence microscopy (see Discussion). However, we have restricted our use of the software to neurons stained using biocytin and a dark reaction product.

**Figure 1:**
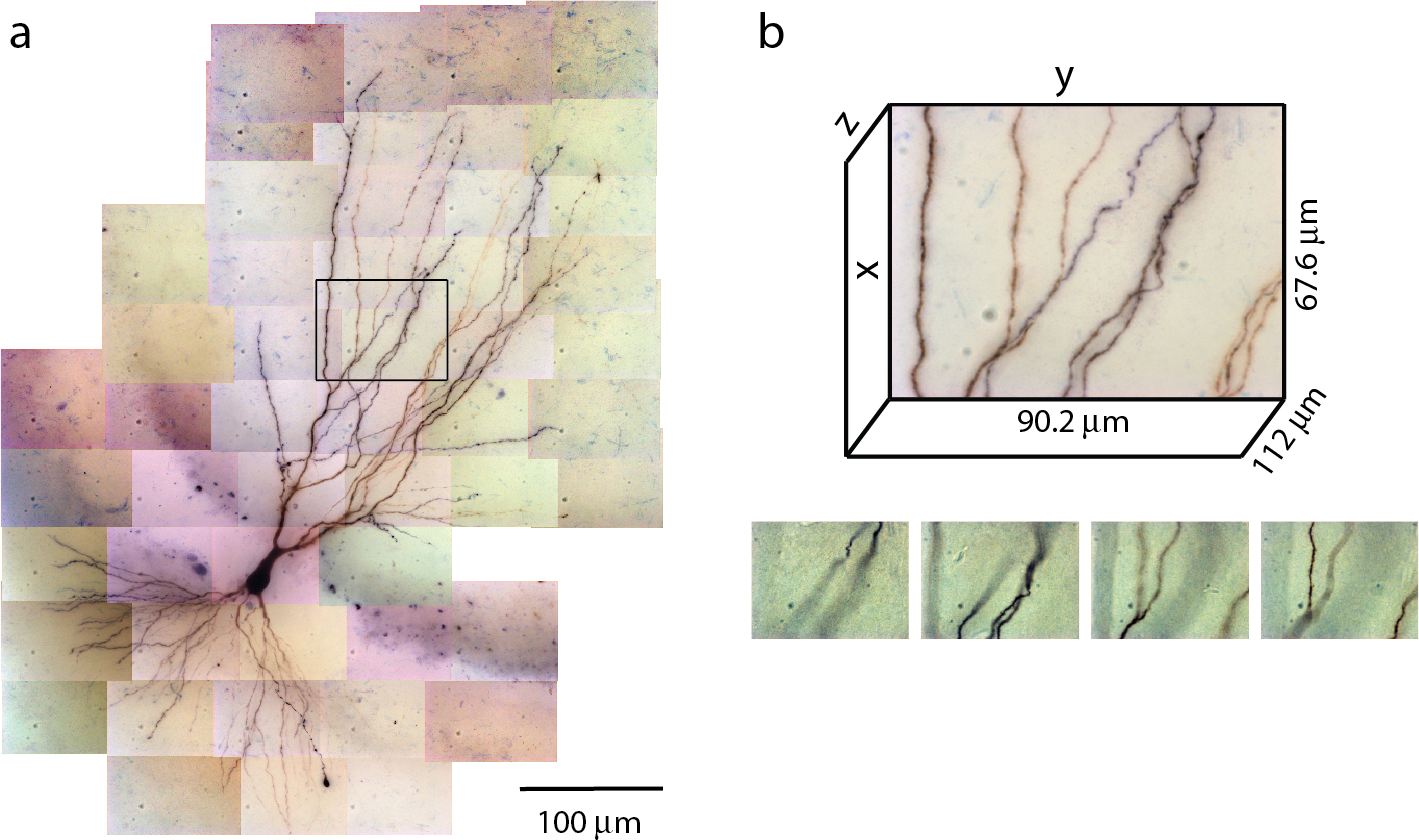
Tiles of tiff stacks covering the entire neuron. a. A mouse CA3 neuron imaged at 100X, NA 1.4. There are 51 tiles covering the entire neuron. 2D projection is shown. b. Dimensions of one tile. Each tiff stack consists of 224 planes of images. The distance between successive image planes is 0.5 *µ*m. Four planes at different depths in the tiff stack indicated by the black rectangle in (a) are shown below.

The number of images required to capture the full three-dimensional (3D) morphology of a neuron depends on its size and the magnification of the microscope objective. The CA3 pyramidal neuron reconstructed here was relatively large and we imaged it using 100X objective (NA 1.4) (Fig. 1). A total of 51 tiles was required, with 224 images per tile (0.5 *µ*m increments through the depth of the slice), thus yielding a total of 11,424 images. This neuron is contained in a volume of *∼* 400 *×* 600 *×* 100 *µ*m^3^ and has a total dendritic length of *∼* 8800 *µ*m. The full imaging process for the CA3 pyramidal neuron took approximately 2 hours on the Zeiss microscope system we used. Faster imaging times (and fewer tiles) can be accomplished using lower magnification objectives, but in our experience 100X provides more accurate estimates of diameters for small-caliber dendrites. During imaging, care was taken to ensure that the microscope settings were optimized to obtain images of all dendrites, including those with the smallest diameter. This resulted in significant background noise, which was removed automatically in a final step of the reconstruction process (see below). We made no attempt to image or reconstruct axons, as these were of finer caliber than dendrites and for many neurons they were difficult to discern beyond a short distance from their origin near the soma.

### Image processing

Because the ZEN blue microscope software provides individual image files in each tile, ShuTu first converts the image files into tiff stacks using the image metadata file (xml) and parsing the file names for depth information. Each tile was imaged successively through the depth of the slice, so no alignment of the images is required to form a stack. As each stack consists of 224 images, about five minutes of CPU time was required for each stack (see Materials and Methods for the system used). The CA3 pyramidal neuron reconstructed here consists of 51 tiles, and creating the stacks required a total of just over four hours. With multiple CPU cores and sufficient memory, ShuTu can automatically distribute the task across multiple cores in parallel, resulting in approximately linear reduction in the real time required to construct the stacks.

After the tiff stacks are created, the tiles need to be stitched to find precise relative positions between the tiles. ShuTu also accomplishes this task in a parallel manner, requiring a similar amount of computational time as construction of the stacks. These two image processing steps are performed in series, but they can be executed sequentially without user intervention. In the case of our example CA3 pyramidal neuron, both of these steps were performed in just a few hours by using multiple CPU cores.

### Automated reconstruction

After image processing, ShuTu produces a draft reconstruction of the neuron using an automatic reconstruction algorithm (Materials and Methods). We devised the algorithm to specifically deal with several challenges posed by the bright-field images of biocytin-filled neurons (Fig. 2). One is background noise (Fig. 2a). While patching a neuron, biocytin can spill out and create blobs in the image stacks. Dirt or dust can be picked up, resulting in structures that look like neurites, especially as color information is not used. Second, during the process of fixing the tissue, thin dendrites can become beaded, with very faint signals between the beads (Fig. 2b). Third, close crossings of adjacent branches require special attention to resolve (Fig. 2c). Fourth, shadows of out-of-focus branches can be as strong as signals from thin dendrites in focus (Fig. 2d), making it hard to trace some dendrites without being fooled by the shadows. These challenges make it difficult to create a perfect reconstruction from automated algorithms. Our algorithm is designed to address many of these issues, but some manual correction is ultimately required. In the following, we outline the steps involved in the algorithm, using the tile shown in Fig. 1b as an example. Technical details of the algorithm are presented in Appendix 2, which should be useful for adjusting the parameters for specific situations encountered by users.

**Figure 2:**
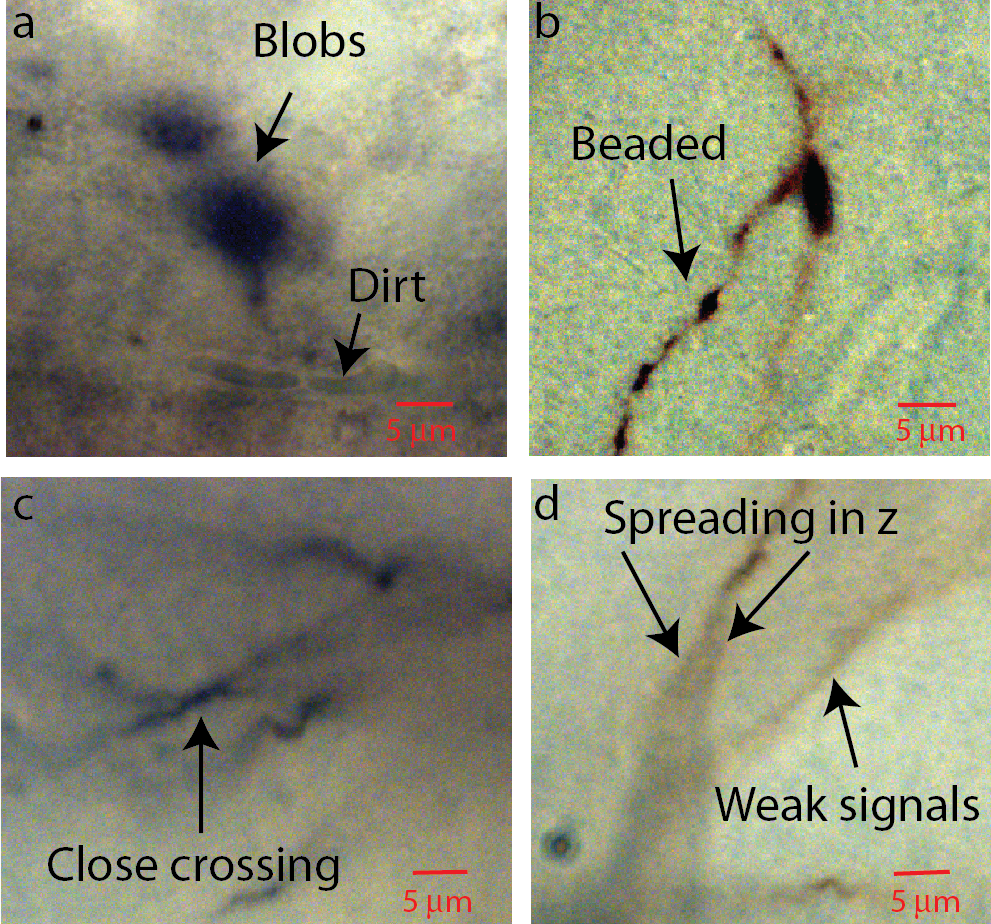
Aspects of bright-field images of biocytin-filled neurons that makes automatic reconstruction challenging. Parts of images in single planes of the tiff stacks are shown. a. Biocytin can be spilled and create spurious signals. Dirt or dust can also add noise. b. Thin branches can be broken into “beads”. c. Close crossing between adjacent neuron branches. d. A branch can cast bifurcating shadows in *z* with darkness level comparable to weak signals from nearby faint branches.

### Conversion to gray scale and 2D projection

The color images are converted into grayscale images, and the pixel intensities are scaled so that the maximum is 1. A minimum intensity projection of the tiff stack is then created, which has the same dimension as a single 2D plane in the stack. The intensity at each pixel is chosen to be that of the darkest pixel among all pixels in the stack having the same *xy* position. This minimum intensity projection reveals all neurites in the tiff stack (Fig. 3a), along with noise from the sources mentioned above. To remove smooth variations due to uneven lighting, the 2D projection is blurred by Gaussian smoothing (Fig. 3b) and subtracted from the original 2D projection (Fig. 3c). Additionally, this process makes faint branches nearly as visible as well-stained ones (Fig. 3d); the inverse peaks corresponding to the branches in the intensity profile have more even heights after the background removal (purple curve) than before (green curve).

**Figure 3:**
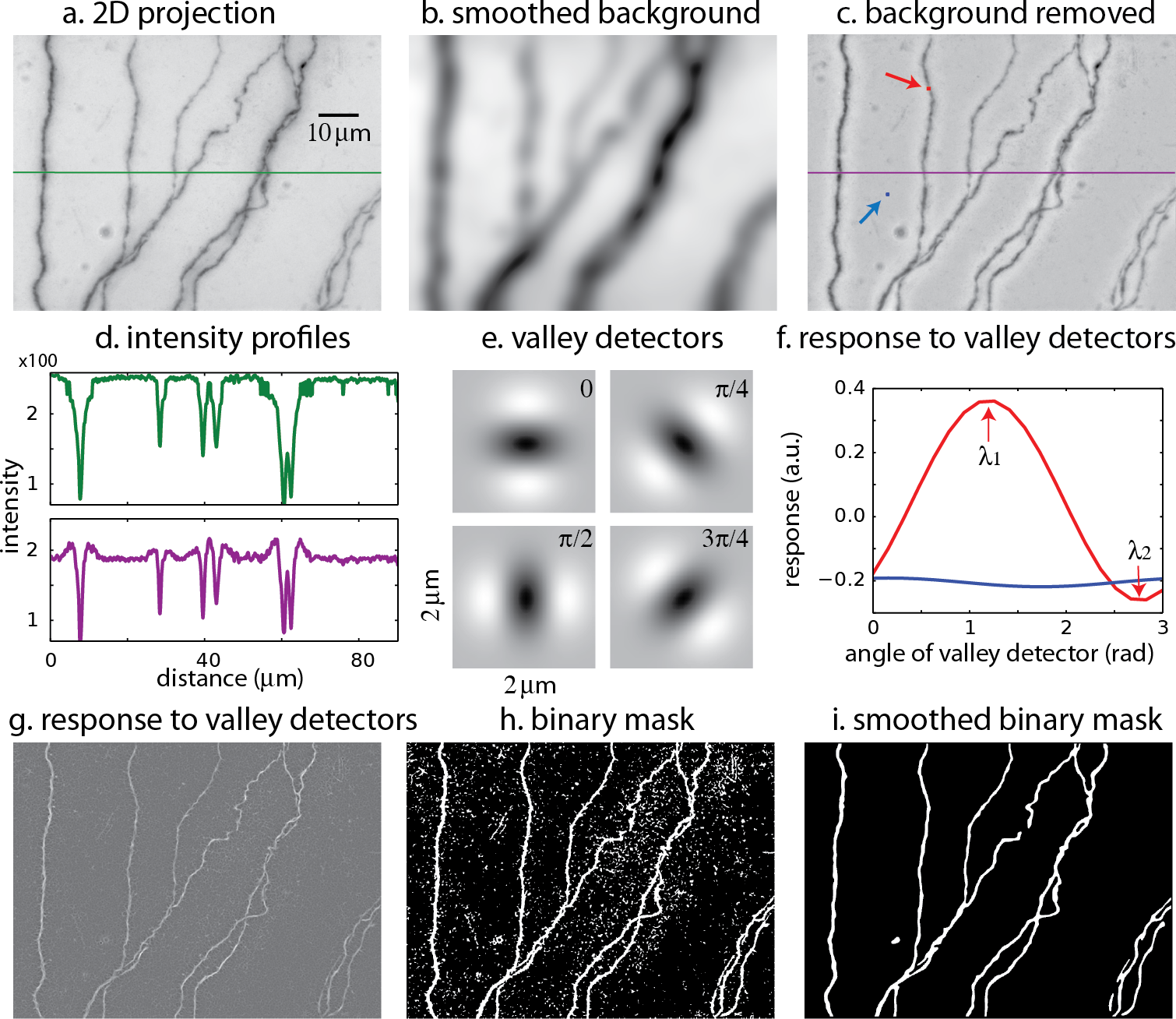
The process of creating a binary mask from 2D projection. a. 2D projection from the image stack. The intensity profile across the green line is shown in (d). b. Smoothed background obtained from Gaussian smoothing of the 2D projection. c. 2D projection after removing the smoothed background. The intensity across the purple line is shown in (d). The red and blue arrows indicate the points to be tested with valley detectors in (f). d. Intensity across the midline in the original 2D projection (green line in (a)) and after removal of the background (purple line in (c)). e. Images of valley detectors at four orientations. f. Responses to valley detectors at varying orientations for two points shown in c (red point, red line; blue point, blue line). *λ*_1_ and *λ*_2_ are the maximum and minimum responses, respectively. g. The maximum responses to the valley detectors (*λ*_1_) for all pixels. h. The binary mask obtained from thresholding *λ*_1_. i. The smoothed binary mask.

### Binary mask

The 2D projection is used to create a mask, which is a binary image with the white pixels indicating the neurites and dark pixels the background (Fig. 3e-i). An accurate mask is crucial for our reconstruction algorithm. Considering the intensity as heights, the neurites in the original 2D projection can be viewed as meandering valleys of dark pixels. To create the mask, we evaluate the possibility that each pixel in the 2D projection belongs to a valley. This is accomplished by comparing the local patch of image centered at the pixel with valley detectors of varying orientations (Fig. 3e) (Frangi et al., 1998). A valley detector is a 2D image consisting of an oriented dark band flanked by two bright bands. The response of the detector is the sum of the products of the corresponding pixels in the detector and the local patch (Fig. 3f). The response has a maximum (*λ*_1_) at one orientation, and a minimum (*λ*_2_) at the orthogonal orientation (Fig. 3f). If the local patch is nearly uniform in intensity, the response is close to zero at all orientations, and *λ*_1_ is small (Fig. 3f, blue curve, which describes the responses at the blue pixel in Fig. 3c). In contrast, if the local patch contains a valley, the maximum response (*λ*_1_) is large and the minimum response (*λ*_2_) is small (Fig. 3f, red curve, at the red pixel in Fig. 3c). If the patch contains a crater corresponding to a blob, *λ*_1_ can be large, but so can *λ*_2_, because there is no previleged orientation. These insights are used to select pixels in valleys but not in blobs or in the background by thresholding *λ*_1_ while also factoring in the difference between *λ*_1_ and *λ*_2_, creating the binary mask (Fig. 3h). The mask is further smoothed to eliminate noisy speckles and rough edges in the boundaries, creating the smoothed mask (Fig. 3i).

### SWC points

The mask is used to place SWC points along the neurites. The SWC points are placed along the centerlines of the binary mask (Fig. 4a). The radii of the SWC points are computed as the shortest distance to the nearest boundaries (Fig. 4b). To determine the depths of the SWC points in the original tiff stack, we dissect the centerlines into segments between end points and/or crossing points. These segments are called ‘xy-paths’ (e.g., Fig. 4a, red arrow). Cutting through the tiff stack while following an xy-path, we create a ‘z-image’ for that segment (Fig. 4c). This z-image contains all pixels in the tiff stack whose *xy* positions lie in the xy-path. The branch whose 2D projection falls on the xy-path manifests as a dark valley in the z-image spanning from the left edge to the right edge (Fig. 4c). ShuTu finds the line through the dark valley (red dotted line in Fig. 4c), from which the depths of the neurites (and the SWC points) are determined. The distance between successive SWC points is set to roughly the sum of their radii. The distance is made shorter when the radii changes rapidly along the centerlines to reflect large changes in short distances in the dendritic morphology.

**Figure 4:**
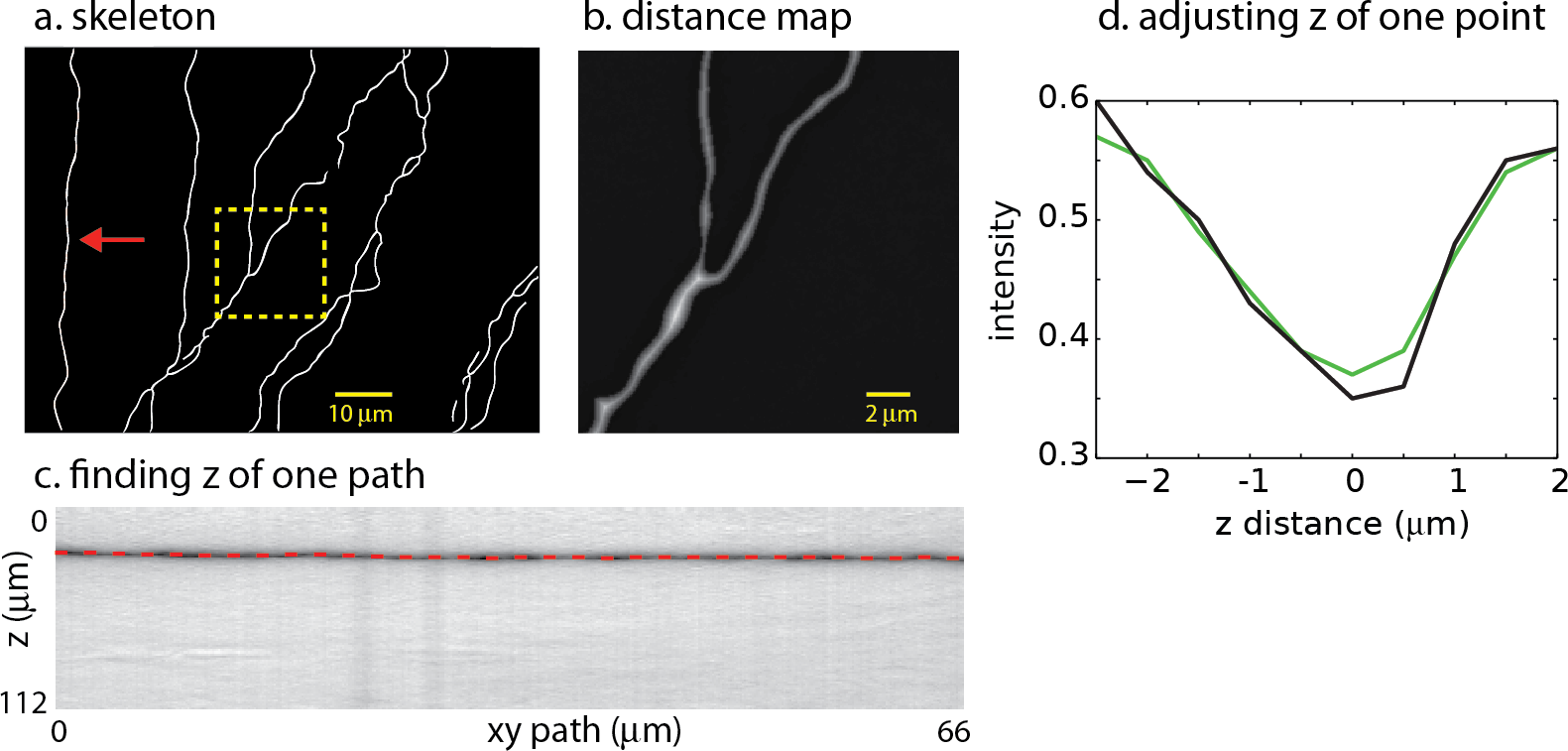
Creating SWC structure from the mask. a. The skeleton obtained by thinning the mask. b. Distance map computed from the mask. The square region highlighted in (a) is shown. The brightness of each pixel is proportional to the distance to the nearest boundary in the mask. c. Finding the depth of the path. The image is constructed by cutting through the stack in *z*-dimension following the *xy* path indicated by the arrow in (a). The dark band is the neurite along the path. The dotted line is the depth (*z*) computed using the left-right shortest path algorithm. d. The depth of a candidate SWC point is further adjusted using the intensity profile in *z* at the *xy* position of the candidate point (black line). The point of minimum intensity in the smoothed profile (green line) is set as the depth of the candidate point. The *z* distance in the graph is relative to the *z* position of the end point.

Invalid SWC points are automatically removed (see below regarding validity of SWC points), and the *z* of a valid SWC point is further adjusted to the nearby depth of minimum of intensity in *z*-dimension (Fig. 4d). Adjacent SWC points along one xy-path are connected. If the removal creates a large distance between two consecutive SWC points, they are not connected. Biologically, sharp turns in neurites are rare. Therefore, to safeguard against possible errors, we do not connect SWC points if doing so creates sharp angles in consecutive lines of connections. To avoid connecting branches far away in depth, SWC points are not connected if the difference in *z* is too large. These decisions depend on parameters set by the user (Appendix 2).

### Validity of SWC points

In some cases, the xy-paths from the centerlines of the binary mask are incorrect. For example, nearby branches can be merged in the mask. Checking the validity of the SWC points is thus crucial for eliminating mistakes. To do so, we take a square patch of the image centered at an SWC point in the plane of the point’s depth. The size of the patch is set to 4 times the radius of the SWC point or 4 *µ*m, whichever is greater. To reduce the possibility that a tilt in the intensity across the patch might interfere with the check, we subtract a linear fit to the the intensity and scale the result to the original intensity range. We then create intensity profiles in eight directions centered at the SWC point (Fig. 5a). For each profile, we look for a significant inverse peak after smoothing the profile (Fig. 5b-c). The significance is checked against the baseline and fluctuations in the intensity. The baseline is set to the top 20% intensity value in the patch, and a parameter *σ* = 0.03 is used to characterize the fluctuations. A threshold, set to the half point between the maximum and minimum of the smoothed profile (Fig. 5b, dotted gray line), is used to judge whether the smoothed profile has two flanks. Another threshold, set to the baseline minus 2*σ*, is used to judge whether the inverse peak is deep enough (Fig. 5b, gray line). If both criteria are met, the profile is judged to have a significant inverse peak. The width of the inverse peak is the distance between the steepest descending point and the steepest ascending point of the peak, identified by the derivatives of the smooth profile (Fig. 5e-f).

**Figure 5:**
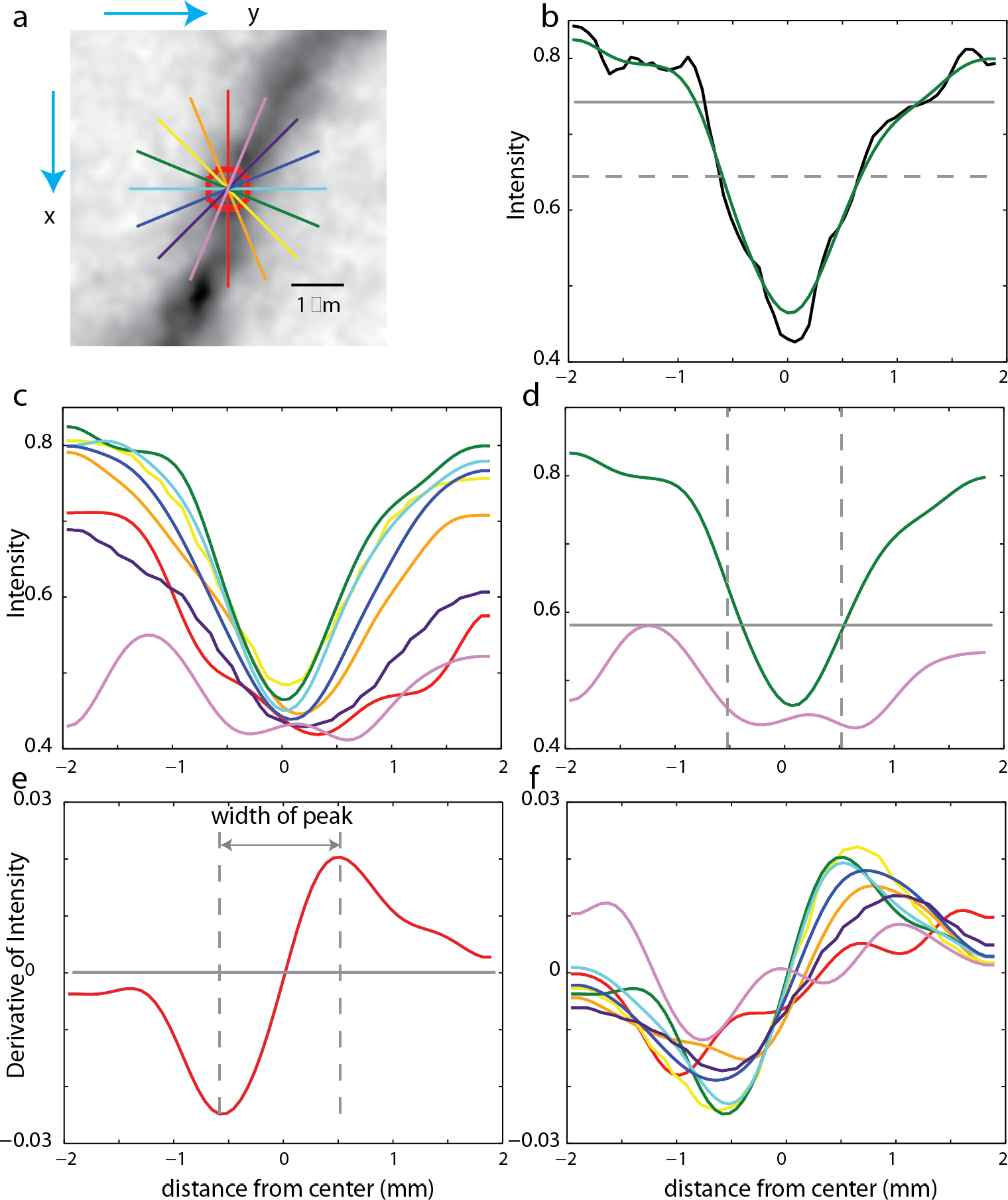
Checking the validity of an SWC point. a. A patch of image around an SWC point to be examined. The image is taken from the *z*-plane of the SWC point. Profiles of the intensities along eight directions are taken (straight lines; colors indicate angles). The green line is the profile chosen to adjust the SWC point. The red circle indicates the radius of the SWC point. b. The profile (black, raw; green, smoothed) along the green line in (a). The dotted gray line is the baseline, and the solid gray line is the threshold. An inverse peak is judged valid if the flanks of the smoothed profile go above the baseline, and the minimum value goes below the threshold. c. Smoothed profiles at all eight directions. d. The chosen profile (green) and the profile at the orthogonal direction (violet). The vertical lines are at the half radius points. Note that the center of the profiles are slightly shifted compared to those in (c). For the SWC to be valid, the minimum intensity of the profile at the orthogonal direction must be below the threshold (gray line) within the vertical lines. e. Smoothed derivative of the smoothed profile in (b). The vertical lines indicate the local maxima of the derivatives. The distance between the vertical lines is the width of the peak. f. Smoothed derivatives of the profiles for all eight directions. The profile with the minimum width is chosen.

If none of the profiles have a significant inverse peak, the SWC point is invalid. Otherwise, we chose the profile with the minimum width among the valid ones. In some cases, an SWC point can be at the edge of thick dendrite or soma (see below). To eliminate them, we check wether the intensity with the half radius of the SWC point is low enough (Fig. 5d). Specifically, we check that the intensity values of the smoothed profile (Fig. 5d, violet curve) orthogonal to the chosen profile (Fig. 5d, green curve) within the half radius (Fig. 5d, dotted vertical lines) is smaller than a threshold. This threshold is set to the maximum of the chosen profile within the range plus *σ*. If not dark enough, the SWC point is invalid.

If the SWC point passes the validity test, we set its radius to the half width of the inverse peak in the chosen profile. Its *xy* position is adjusted to that of the inverse peak, and *z* position is adjusted to the depth of the nearby intensity minimum in *z* (Fig. 4d). To ensure that this adjusting process converges, we adjust each SWC point three times iteratively. If the final *xy* position shifts from the original position more than twice of the original radius, we mark the SWC point invalid since it is most likely created erroneously. Finally, if the final radius of the SWC is smaller than 0.2 *µ*m or larger than 10 *µ*m, the SWC is most likely due to noise and is marked invalid.

### Mark pixels occupied

As the SWC points are created, we mark pixels in the tiff stack in the vicinity of the SWC points as occupied (Fig. 6). Before creating a new SWC point, we check whether its center point is marked as occupied. If so, no SWC point is created. This avoids creating redundant SWC points for the same piece of dendritic branch.

**Figure 6:**
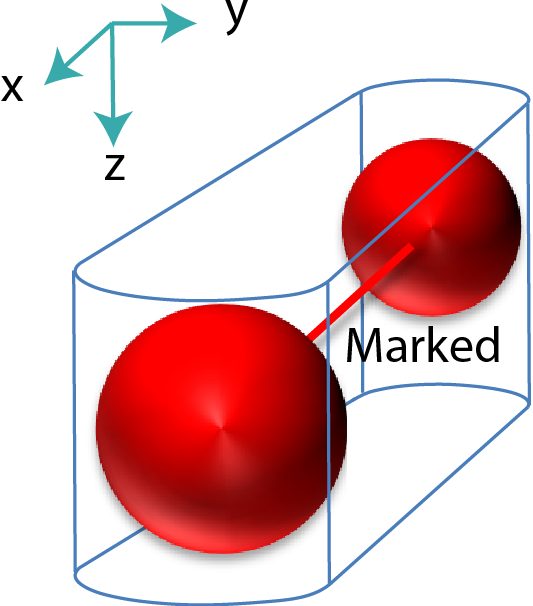
Mark pixels in tiff stack occupied. The pixels around two connected SWC points (red spheres), formed by two half cylinders and a trapezoidal prism, are marked as occupied.

### Thick dendrites and soma

The widths of dendrites can vary by as much as five times from the thin terminal dendrites to the thick apical dendrite near the soma. The thick dendrites and the soma can be missing from the binary mask, which is created with the valley detector tuned for detecting thin dendrites. Instead, only edges of the thick dendrite and soma are captured in the mask, leading to invalid SWC points that are eliminated. To solve this issue, we specifically detect the presence of thick dendrites and soma. The thick dendrites and soma are typically well-stained and show up as darkest parts in the 2D projection. We use this fact to decide whether there are thick dendrites and soma that are not been covered by existing SWC points. If the lowest intensities in the pixels covered by the existing SWC points are brighter than the lowest intensities in the 2D projection, we decide that the binary mask missed the soma or thick dendrite. We create a binary mask on a 2D projection, excluding pixels around the existing SWC points. New SWC points are added based on this mask.

### Extending SWC points in 3D

The SWC structure created with 2D projections can contain errors. Typically the binary masks can be incomplete or wrong in some parts due to weak signals, occlusions produced by brach crossing, or mergers of closely parallel branches. This leads to gaps in the SWC structure representing continuous dendritic branches. To bridge these gaps, we extend the SWC points in 3D (the tiff stack) from the end points in the SWC structure.

To minimize the interference from noise, we first delete isolated SWC points that are not connected to any other SWC points. We then mark pixels near the existing SWC points occupied (Fig. 7a, red circles) to ensure that the extension does not create duplicated SWC points.

**Figure 7:**
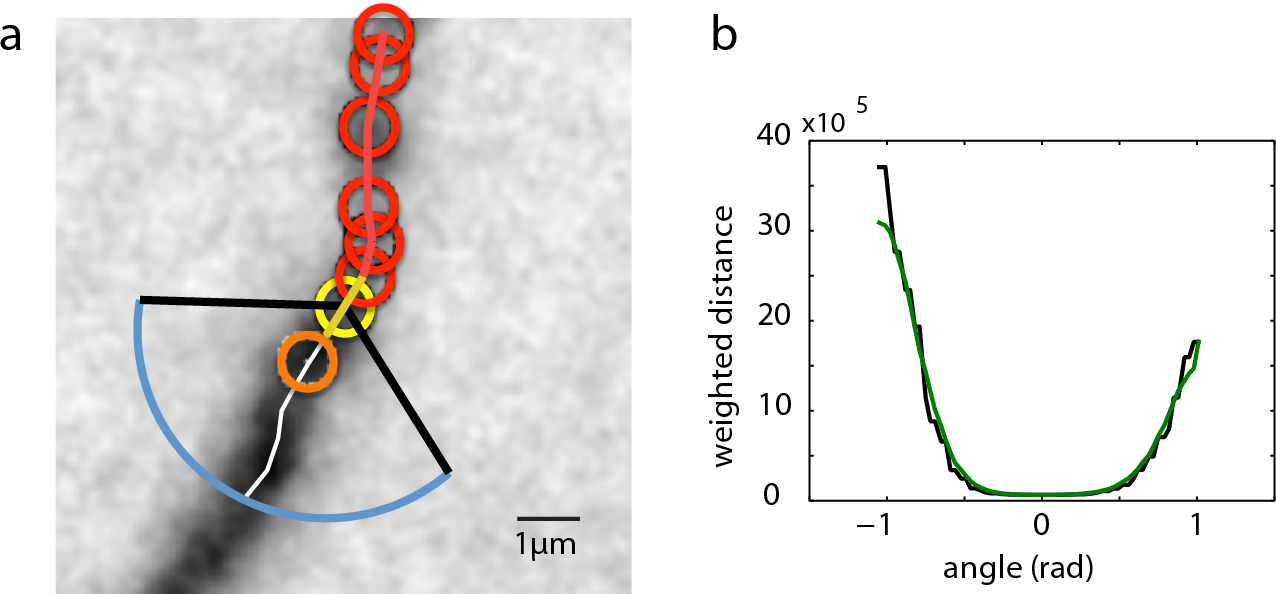
Extending SWC structure in 3D. a. The candidate point for extension is searched in the plane of an end SWC point (yellow circle). Red circles are the SWC points that are connected to the end point. The search is done along a shortest distance path running through the neurite (white line). This path is determined by building the intensity-weighted shortest distance profile along an arc (blue line) enclosed by two black lines. The valid SWC point closest to the end point is selected as the new SWC point (orange circle). b. The profile of the shortest distances along the arc (blue line in (a)). The green line is the smoothed version. The angle is measured relative to the line connecting the end point to its connected SWC point (yellow line in (a)). The minimum position in the smoothed profile is selected as the starting point of the shortest path shown in (a).

From an end SWC point (Fig. 7a, yellow circle), we search for the next candidate SWC point. We draw an arc of radius 3 *µ*m or twice the radius of the end point, whichever is greater, in the plane of the end point (Fig. 7a, blue arc). The arc spans from *−π/*3 to *π/*3 (Fig. 7a, black lines) relative to the line from the end point to its connected SWC point (Fig. 7a, yellow line). The shortest intensity-weighted distances from the points on the arc to the end point are computed (Fig. 7b, black line). With the smoothed profile of the distances, the point with the minimum distance is selected (Fig. 7b, green line), and the shortest distance path from this point to the end point is found, which should follow along the neurite (Fig. 7a, white line). The depth of the neurite is found using the xy-path technique (Fig. 4c) along the shortest distance path.

The candidate SWC point for extension (Fig. 7a, orange circle) is placed on the shortest distance path, starting from the end point marching towards the arc. We test the validity of the candidate SWC point, during which the *xy* position and radius are adjusted. To cover weak branches, the test is made less stringent by accepting shallower inverse peaks in the intensity profiles used for the test (Fig. 5b-c). If accepted, the extension process continues from the new SWC point as the end point. If the candidate point is marked occupied, the extension stops, and the possibility of connecting the end point to the existing SWC points that marked the occupation are evaluated (see below). The extension stops if the test fails for all points along the shortest distance path.

### Connecting broken segments

After extending SWC points in 3D, a continuous branch can still be represented with broken segments of SWC points, especially if the underlying signal is broken or there are closely crossing branches (e.g., Fig. 2b,c). We connect these segments with heuristic rules based on the distances between the end points, in order to recover the branch continuity (Appendix 2). After connecting the end points, the SWC structure for the tiff stack is complete.

The results for our example tiff stack are shown in Fig. 8, in which the SWC points are overlaid with the underlying image, and in Fig. 9, in which the SWC structure is shown in four different view angles in 3D to reveal more details. For this particular tiff stack, the automated reconstruction is mostly accurate, except that an elongated piece of dirt is mistaken as a neurite, and a close crossing of two branches is incorrectly connected. These errors need to be corrected manually (see below).

**Figure 8:**
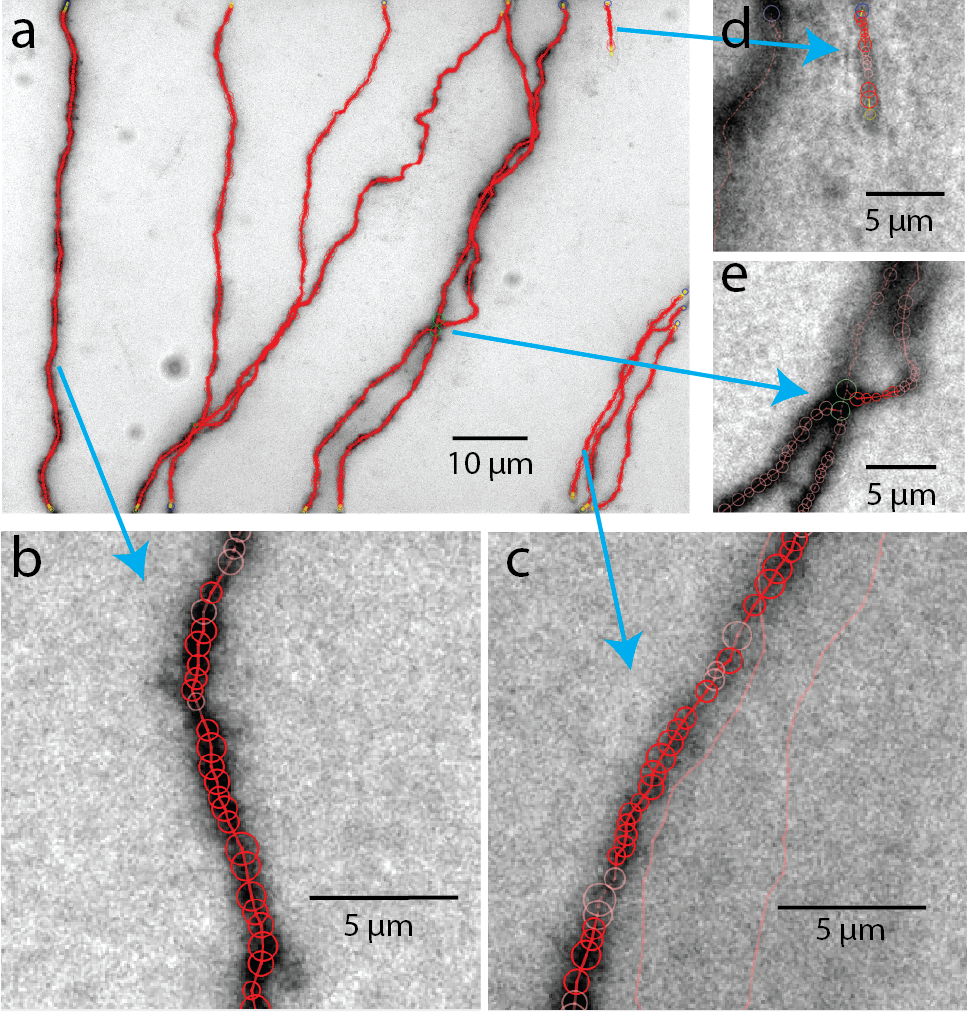
The SWC structure overlaid on the image. a. The SWC structure projected on to the image of 2D projection. Red circles are SWC points. Connections between them are indicated with red lines. b-d. SWC structure overlaid at the specific planes in the tiff stack, zoomed in to show more details. Arrows indicate the corresponding regions in the 2D projection.

**Figure 9:**
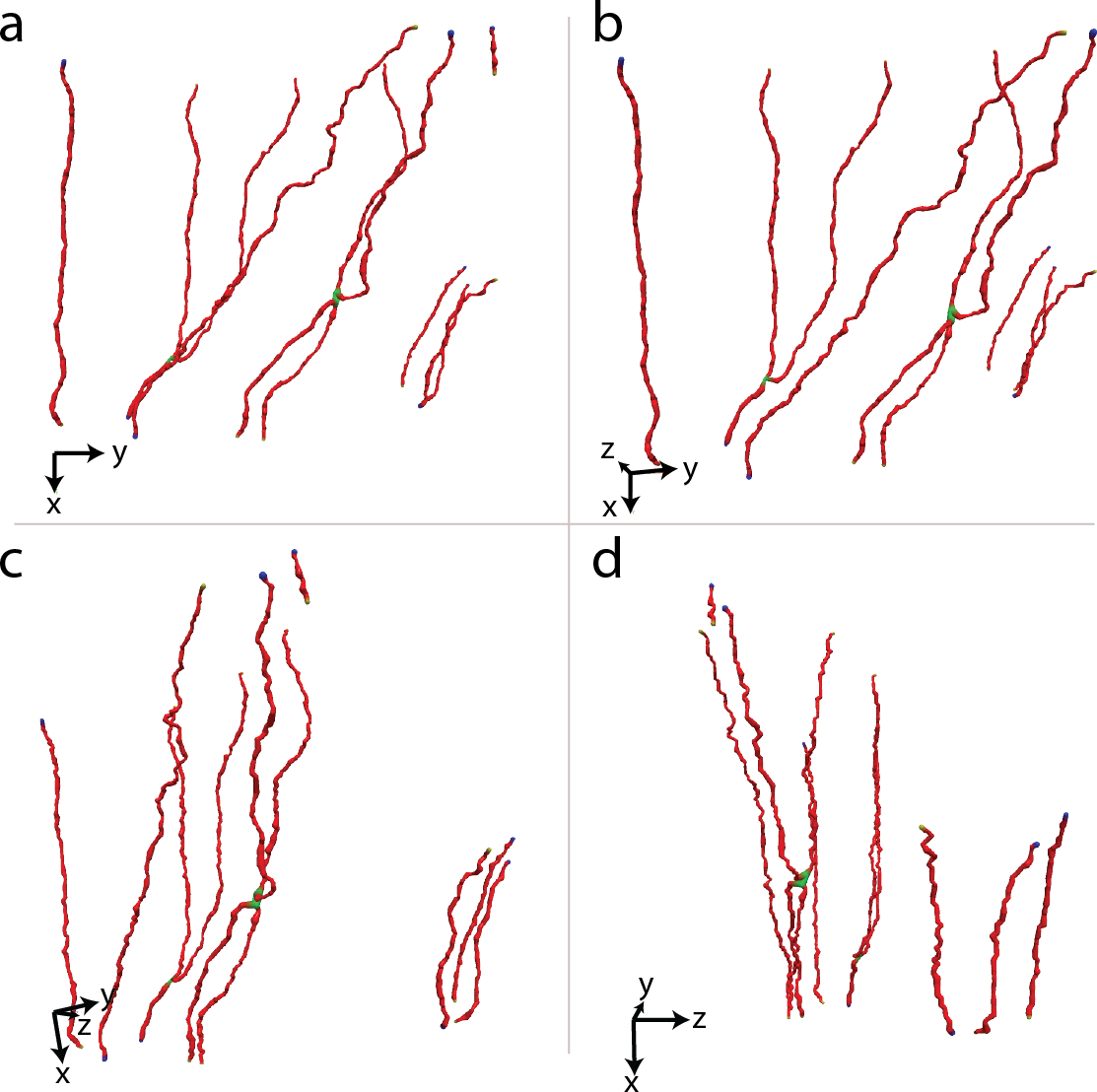
The SWC structure viewed from four different angles to reveal the 3D structure. The viewing angles are indicated with the directions of *xyz* coordinates. The view angle in (a) corresponds to the 2D projection view.

### Subdivision in *z*

The 2D projection can be complicated when there are many branches in one tiff stack, which often leads to missed branches due to occlusions. One way of mitigating this problem is to divide the tiff stack in *z* into several slabs with equal heights in *z*. SWC points are created separately for each slab as described above, and then combined for the entire stack. The extension from the end points is done with the entire stack. When branches extend across the boundaries between subdivisions of tiff stacks, they are automatically connected by extension from the end points, as described above.

ShuTu allows the user to decide how many subdivisions are necessary based on the complexity of the morphology and the thickness of the tiff stacks. The user should keep in mind that a large number of subdivision slows down the automated tracing. In our example neuron, we divided all tiff stacks into eight slabs.

### Combining SWCs

The SWCs of individual tiles of tiff stacks are combined to form the SWC of the entire neuron. The positions of SWCs are shifted based on the relative coordinates obtained in the stitching process. The SWC points of individual stacks are read in sequentially. To void duplicated SWC points in the overlapping regions of adjacent stacks, pixels near the SWC points that are already read in are marked occupied. If the position of SWC points are at the marked pixels, they are deleted. After reading in the SWC points of all stacks, we extend the end points and connect them if they are nearby. Isolated short branches (< 20 *µ*m) and small protrusions (< 5 SWC points) from main branches are deleted to reduce noise in the SWC structure. The resulting SWC structure for the example neuron is shown in Fig. 10a-d.

**Figure 10:**
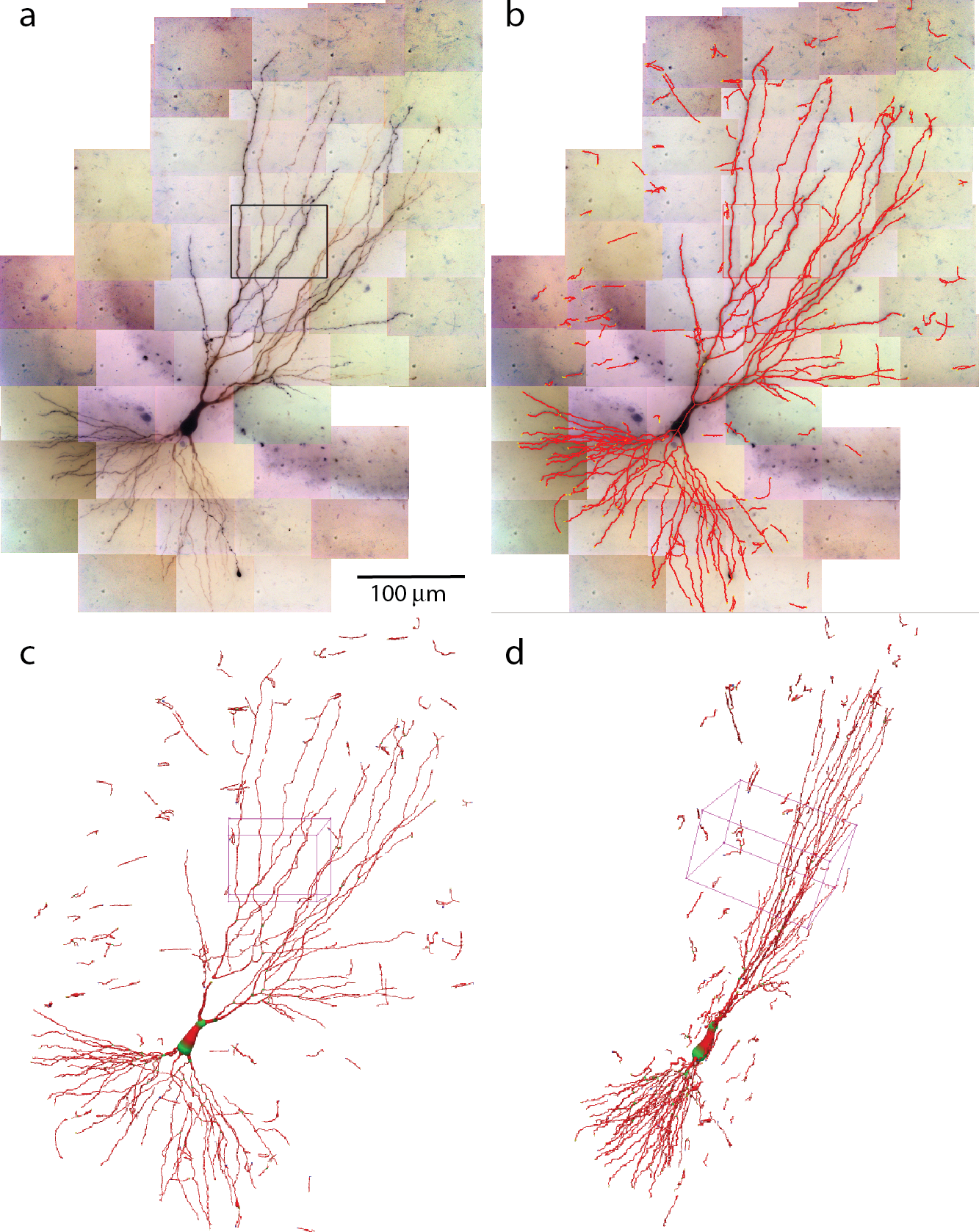
Combing SWCs from the stacks for the entire neuron. a. 2D projection of the entire neuron. Individual tiff stacks are stitched together to obtain their relative coordinates. The stack in Fig.9 is highlighted with a black rectangle. b. The SWC of the entire neuron is obtained by combining the SWCs of individual tiff stacks. The SWC points are overlaid onto the 2D projection. c. 3D view of the SWC structure. The 3D box corresponds to the highlighted stack in (a). d. The 3D view from a different angle.

The entire process of automated reconstruction of the example neuron took about four hours on our basic desktop system using three CPU cores (see Materials and Methods for the system specs). With more powerful computers, the time can be further reduced approximately linearly with the number of CPU cores used.

### Manual editing and error correction

The SWC structure created by the automatic algorithm requires editing, such as removing noise, tracing thin or faint dendrites, connecting ends, and correcting mistakes in the radii and positions of the SWC points and in the connections between them. We have designed ShuTu to make these operations easy for the user. In this section we highlight a number of editing techniques.

### Inspecting the reconstruction

The SWC structure can be examined in three ways: Tile Manager, Stack View, and 3D View (Fig. 11). In Tile Manager, the SWC structure is overlaid with 2D projection of the entire neuron (Fig. 11a). In this view, it is easy to identify missing, discontinuous, or incorrectly connected branches.

**Figure 11:**
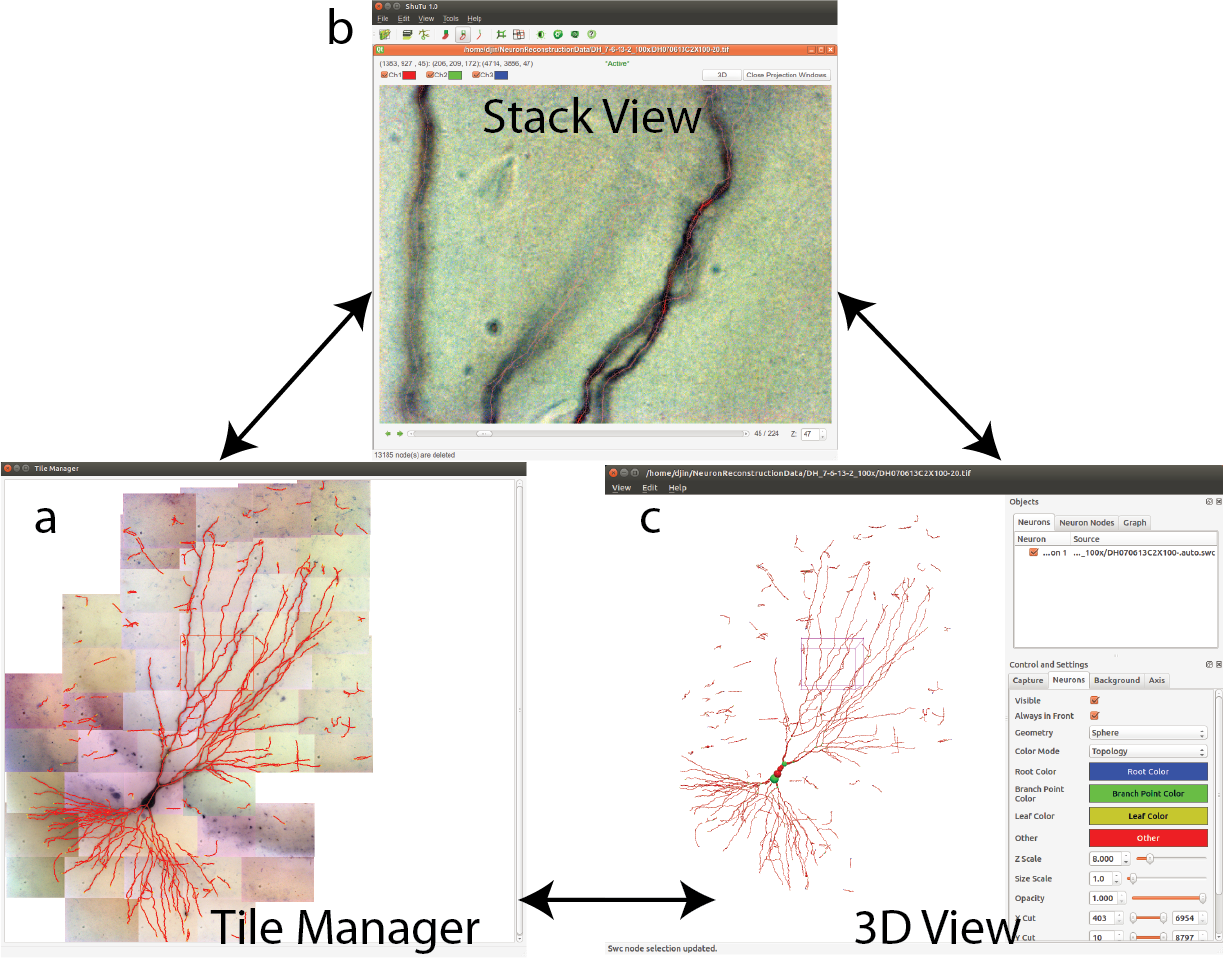
Three views for examining the SWC structure. a. Tile manager. 2D projection of the stitched stacks is superimposed with the 2D projection of the SWC structure. b. Stack view. One stack is loaded, with the SWC points in the stack overlaid onto the image. The 2D projection view of the stack can be created within this window. c. SWC view. The 3D structure can be viewed from different angles and edited.

Double clicking on one tile in Tile Manager loads the tiff stack into Stack View (Fig. 11b), in which the SWC structure is overlaid with the image. The radii, depths and connectivity of the SWC points can be examined in detail by scrolling up and down through the *z* dimension of the tiff stack.

From Stack View, a 2D projection can be created by clicking on Make Projection button (Fig. 12). There is an option to subdivide the stack into multiple slabs in *z*, in which case separate 2D projections are created. Subdivision is useful when the branching patterns are complicated. Mistakes in the reconstruction can be easily spotted in Projection View, including missed branches, broken points, incorrect connections, and inclusion of noise (Fig. 12). Incorrect positions and diameters for the SWC points are easy to identify as well.

**Figure 12:**
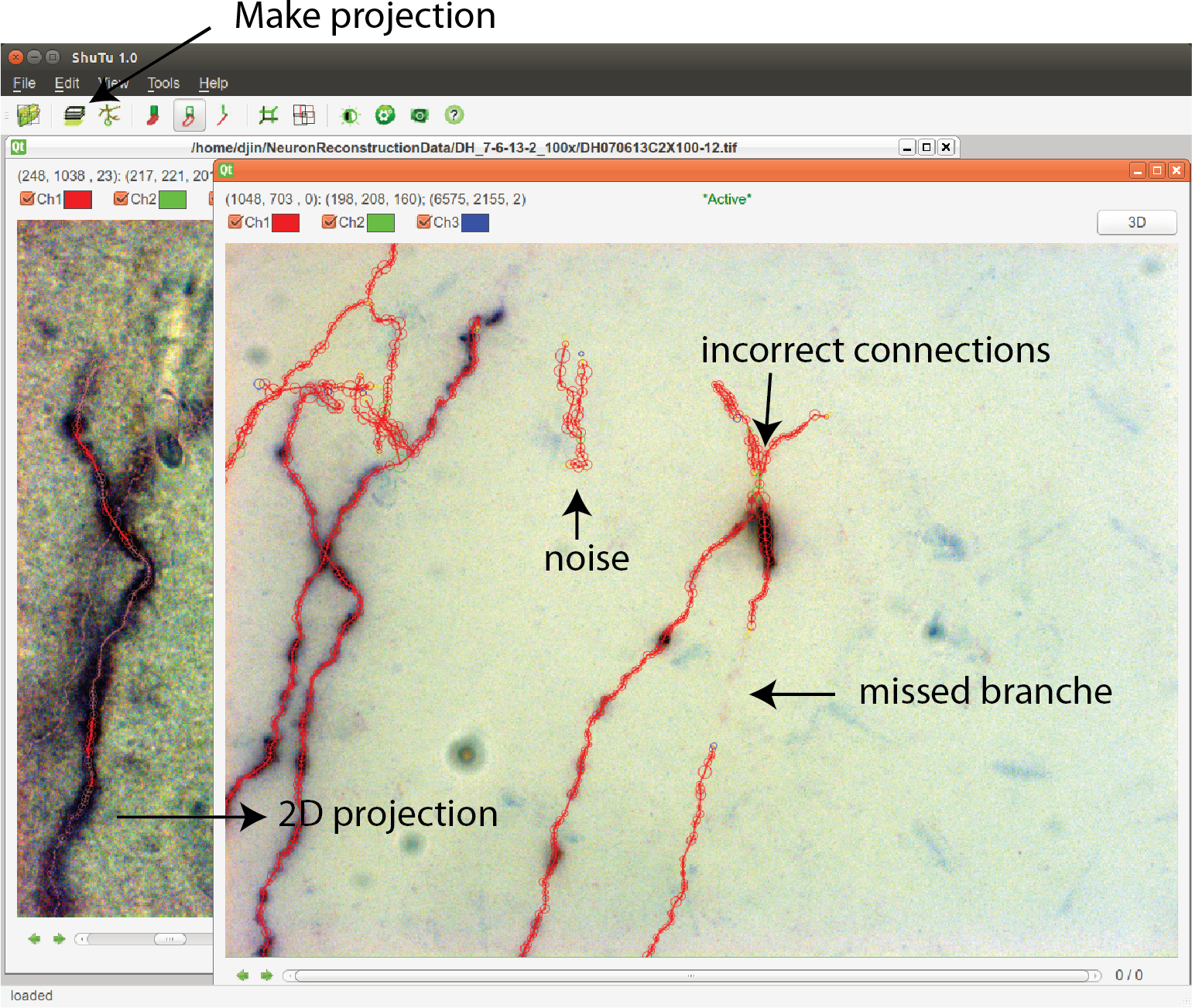
In the Stack View, clicking on the Make Projection button creates the 2D projection of the tiff stack and the SWC structure. It is easy to spot mistakes in this view. SWC points can be removed and their properties changed. The connections between SWC points can be modified. Selecting one SWC point and pressing z locates the points in the Stack View for further examination and modification.

In 3D View, the SWC structure can be rotated and shifted in order to reveal incorrect connections, especially large jumps in *z*, which can be obscure in other views.

Editing can be done in Stack View, Projection View, and 3D View. In all cases, after any editing, the SWC structure is updated in all views. A selected point can be deleted or moved and its radius can be modified. A selected point in Projection View or 3D View can also be located in Stack View for further examination and modification using the stiff stack.

### Adding SWC points

In Stack View, SWC points can be added in three ways. The first method is smart extension. The user selects an SWC point on a branch that needs extension, finds a target point on the branch and locates the focus plane in *z*, and then clicks on the target. SWC points will be added along the branch from the selected SWC point to the target point (Fig. 13a). The path is computed with the shortest distance algorithm, and the radii and positions of the SWC points are automatically calculated using the automated algorithm described above. The second method is manual extension. It is the same as the smart extension, except that the only point added is at the target point and its radius needs to be adjusted manually. The third method is mask-to-SWC (Fig. 13b). In Projection View, a mask along a branch is drawn by selecting the start and end points. The path is automatically computed with the shortest-distance algorithm. The mask can also be drawn manually. After the mask is completed, it is converted to SWC points along the branch. The positions and radii of the SWC points are computed automatically.

**Figure 13:**
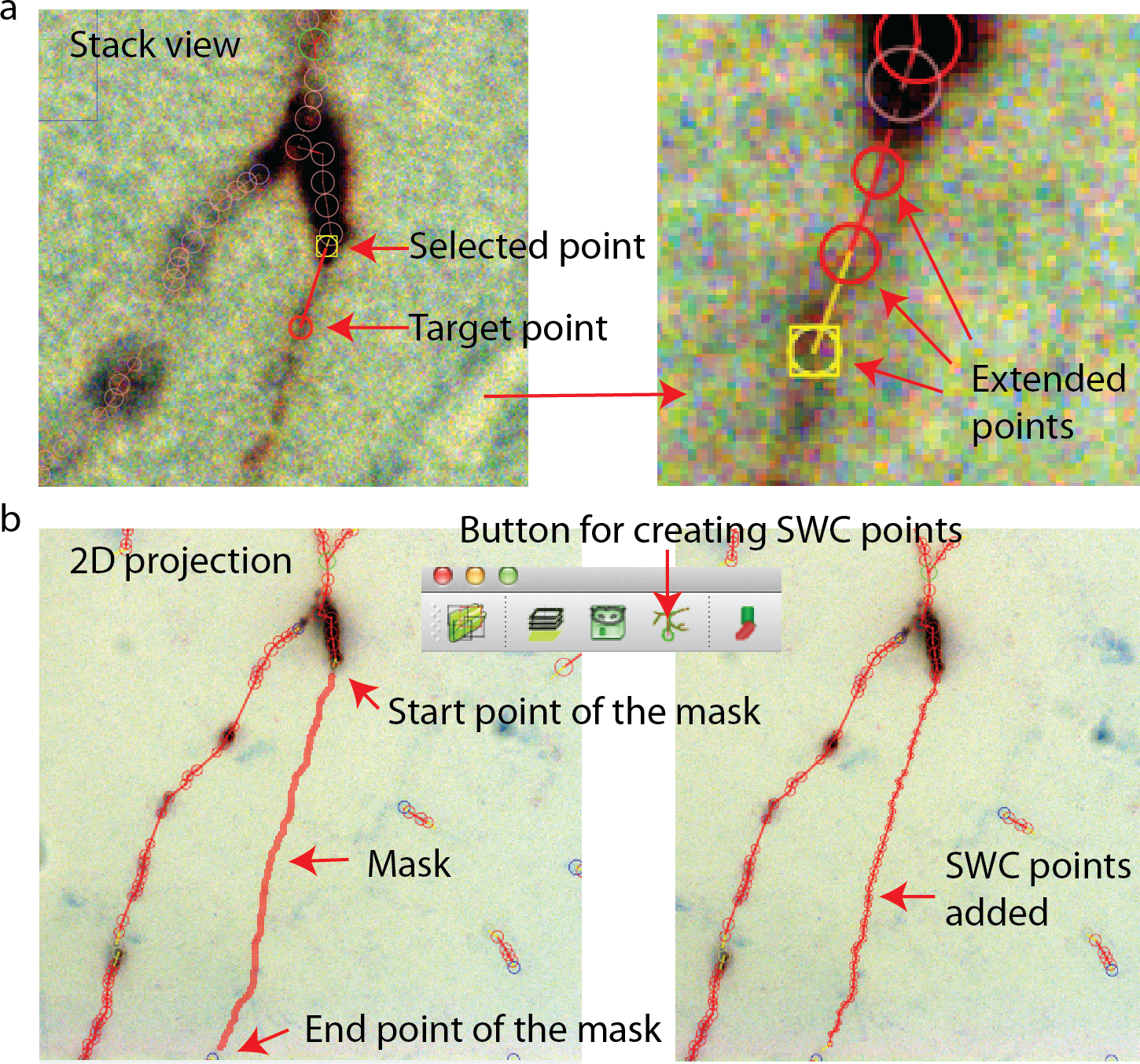
Creating SWC points. a. In Stack View, an SWC point is selected. Find the target point by finding the focus plane of the branch. Clicking on the target point creates SWC points connecting the target point to the selected point along the branch. b. In Projection View, pressing r starts mask creation. Click on the starting point and `Shift-click` on the end point along a branch creates a mask. Clicking on the `Mask → SWC` button creates SWC points along the mask.

These three ways of adding SWC points are complimentary. When the branch to be reconstructed is long, the mask-to-SWC method is efficient. However, it requires that the underlying signal is strong enough, otherwise the computation of the path and the depths can be inaccurate. When the branch to be covered is short, the smart extension method is efficient, although it also requires a relatively strong signal. Manual extension always works.

ShuTu users can reconstruct the entire neuron with one of these three methods. The extension methods can be used after creating a single seed SWC point. However, the process is tedious because the focus plane must be located in every click. The mask-to-SWC method traces branches in 2D projections, and is therefore more efficient.

### Modifying connections

The end points in the SWC structure are highlighted with blue or yellow colors. In some cases, it is necessary to connect nearby points that have been incorrectly identified as end points. This can be done by selecting two end points and connecting them. If the distance between the two points are more than the sum of their radii, SWC points can also be added automatically while bridging the gap. A selected end point can also be automatically connected to its nearest neighbor.

Incorrect connections can be broken after selecting two connected SWC points. The branching points are highlighted with green; these points need to be examined carefully for incorrect connections, especially when branches cross.

All SWC points connected to a selected point can be highlighted (Fig. 14a). This is useful for finding broken connections in the SWC structure. Structures with short total length can also be selected (Fig. 14b), and can be deleted with a single command. This is useful to reduce noise in the automated reconstruction. At the end of the reconstruction, all SWC points that belong to the neuron should be connected. At this point, all remaining points (presumably noise) can be deleted by selecting all connected points in the neuron and deleting all unconnected points.

**Figure 14:**
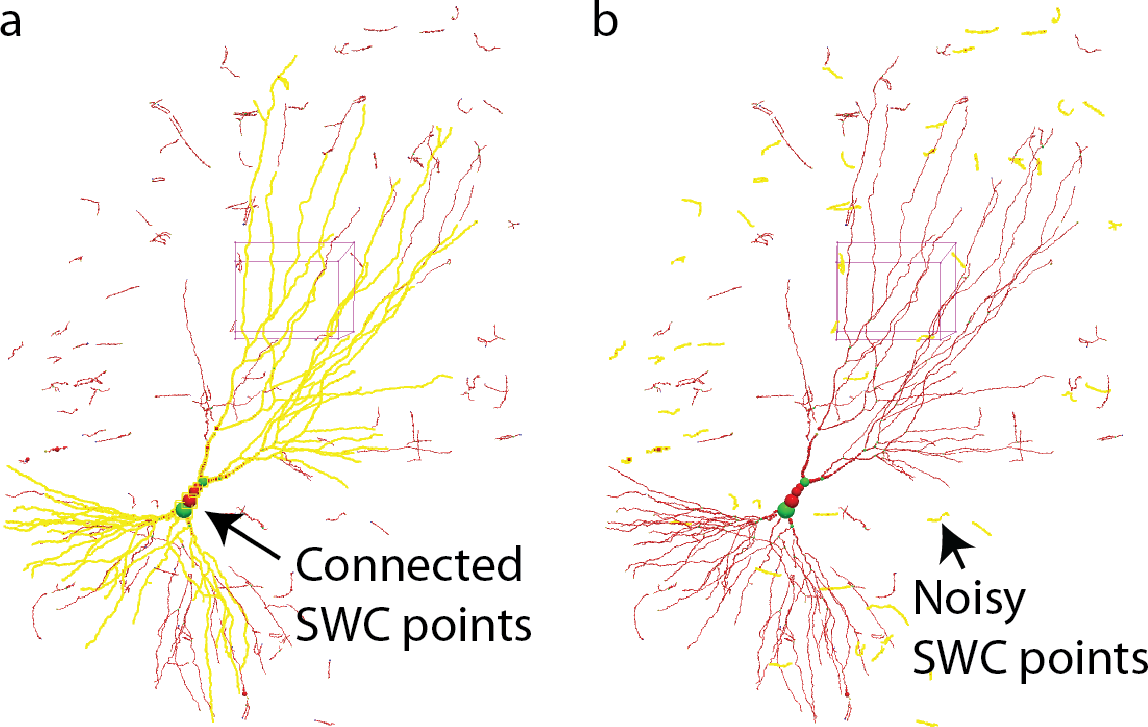
a. In 3D View, selecting one SWC point and pressing h-5 selects all SWC points connected to the selected. This operation is useful for detecting broken connections. b. Pressing h-7 selects all branches with total length smaller than a chosen threshold. This operation is useful for deleting noise.

### Reconstruction efficiency

To quantify the efficiency of reconstructing neurons through the automatic algorithm and manual editing, we counted the number of editing operations (NEO) required for achieving the final reconstructions starting from the one generated by the automatic algorithm. The results for the example neuron are shown in Fig. 15. The SWC points that are added in the editing phase are shown in red, and those from the automatic reconstruction are shown in green (Fig. 15a,b). The added SWC points are about 5% of the total SWC points in the structure. The NEO is 439. Among the editing operations, extensions are dominant. Correcting connection mistakes are sizable as well. The manual time spent in repairing the automatic reconstruction was around 1.5 hours.

**Figure 15:**
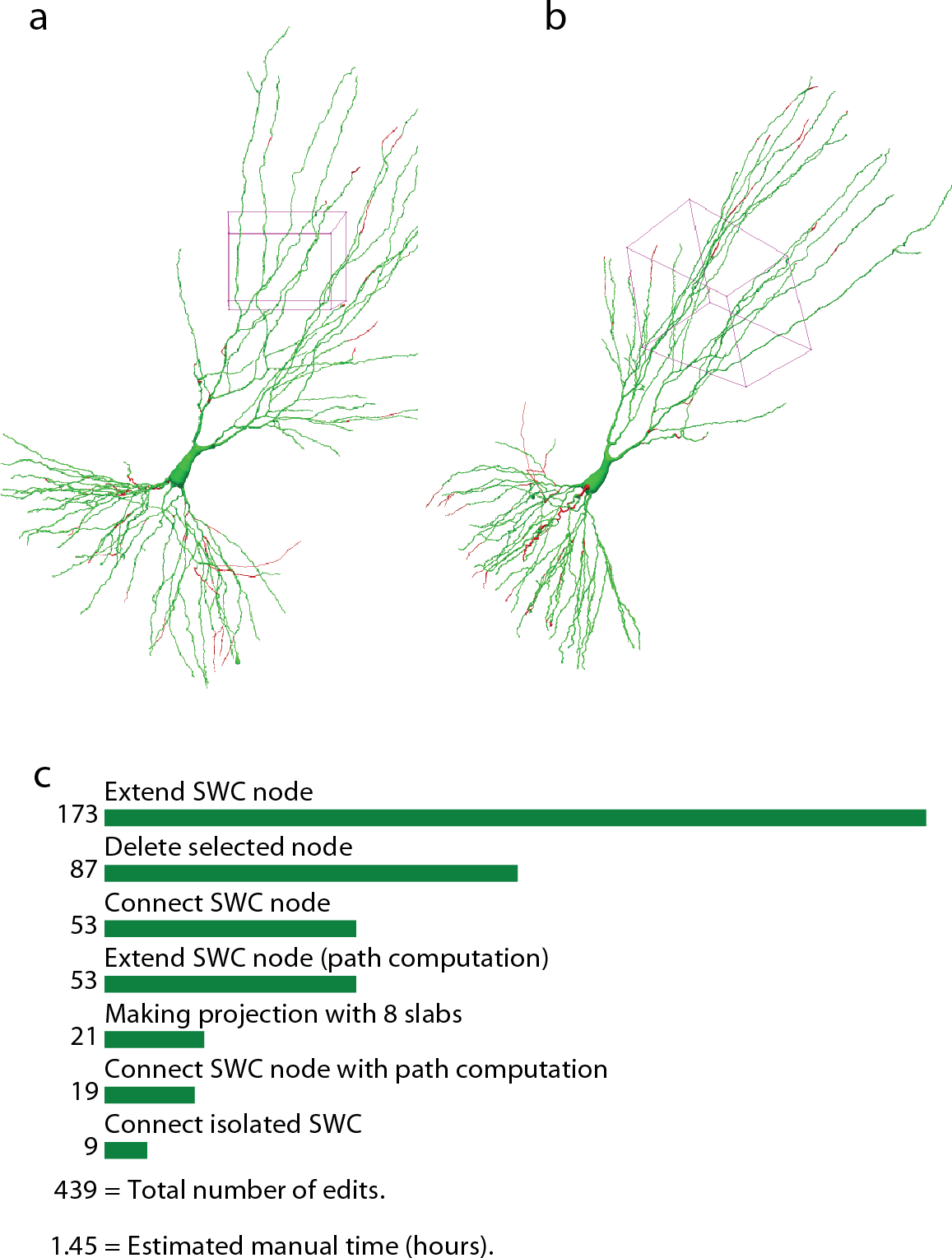
Reconstructed neuron after editing. a,b. Two different views of the reconstructed neurons. The SWC points from the automatic reconstruction are in blue, and those added in the editing process are in red. c. Top operations done in the editing process and the total number of edits.

The efficiency of reconstruction depends on the image quality and the complexity of the neuron morphology. For neurons with sparse processes, the automated reconstruction captures the most of neuronal structure, and manual editing is not intensive. For the example shown in Fig. 16a, which is another mouse CA3 pyramidal neuron, simpler than the one shown in previous figures, the NEO is 88, and the time spent in editing was approximately 20 minutes. In contrast, when the processes are dense, the automated reconstruction contains many misses and mistakes, and manual editing takes more efforts. An example is shown in Fig. 16b, which is a rat CA1 pyramidal neuron; the NEO is 812, and the time spent in editing was approximately 1.9 hours. The complexity of the processes requires more time for examining the appropriate dendritic structures. Another example of a complex neuron is shown in Fig. 16c, which is a mouse Purkinje cell imaged with confocal microscope. The increased complexity decreases the quality of automated reconstruction, and the NOE is 1190, leading to approximately 2.4 hours of editing.

**Figure 16:**
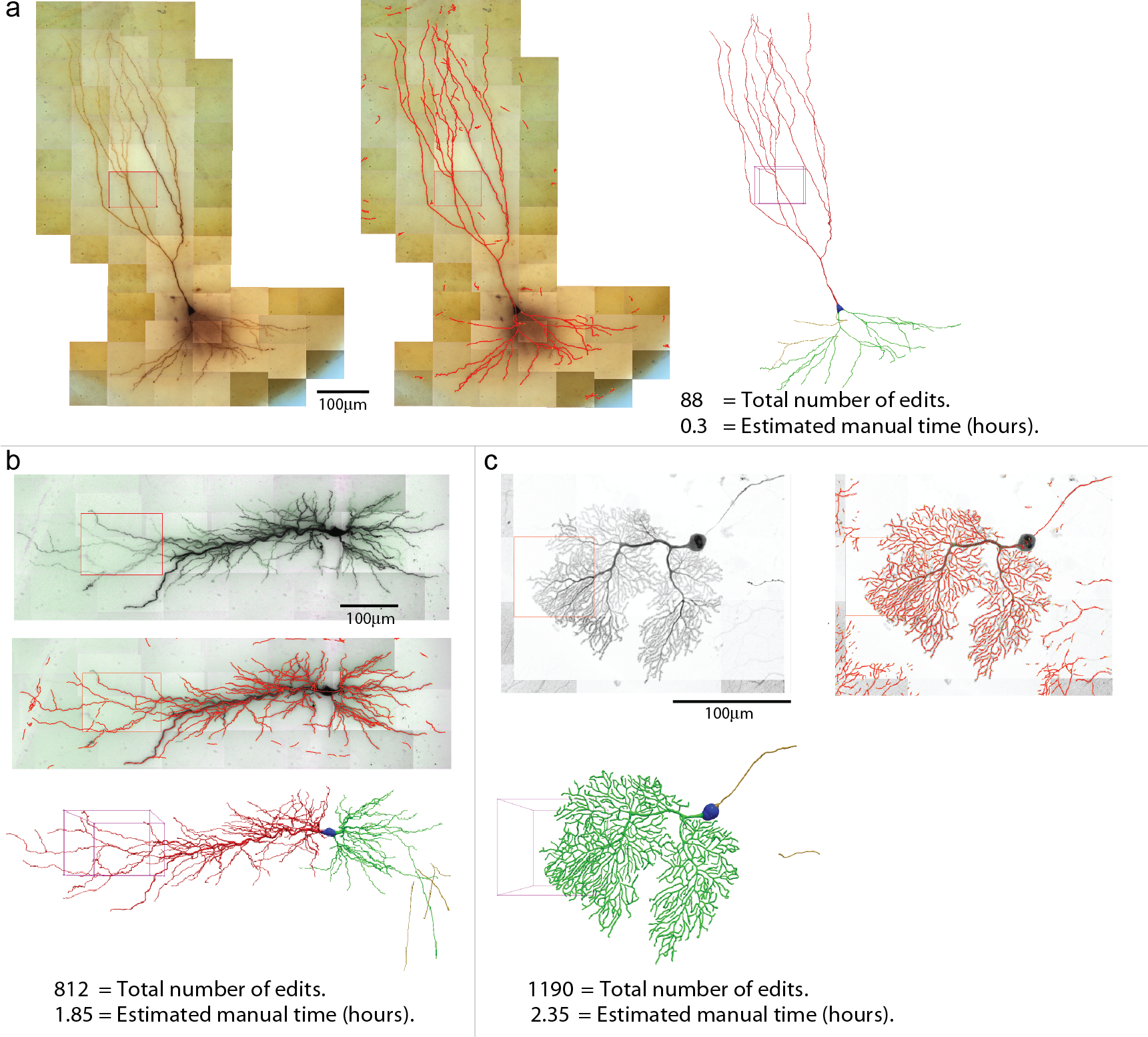
Three examples of reconstructions. Shown for each neuron are the 2D projection of the images, automated reconstruction on top of the 2D projection, and the final reconstruction (blue, soma; red, apical dendrite; green, basal dendrite; gold, axon). a. A pyramidal neuron in the mouse CA3 region imaged at 100X (biocytin). b. A pyramidal neuron in the rat CA1 region imaged at 63X (biocytin). c. A mouse purkinje cell imaged at 63X with confocal microscope.

## Discussion

We have demonstrated how ShuTu can be used to reconstruct neuron morphology by converting microscope images to SWC structures. Our goal is to provide a practical system that can be readily implemented and used in labs who need accurate dendritic reconstructions of neurons, often that have been studied and stained following recordings with patch-clamp electrodes. As an open-source software package, it can be continuously improved by the community. We have also provided raw images of the example neurons (Jin, 2019), which should be useful for testing and improving the software.

A major aim of our software package is to minimize human labor in reconstructing neurons. We introduced editing functions in ShuTu to improve the efficiency of editing, and implemented a method for counting the number of editing operations (NEO) as the measure of the success of automatic reconstruction. The algorithm for reconstruction and editing functions were developed with the goal of optimizing the automated reconstruction and reducing NEO. For example, our automatic reconstruction algorithm can be aggressive in finding neurites, including faint ones, despite the fact that this may lead to inclusion of more noise in the reconstruction. During the editing phase, this noise can be easily eliminated once the SWC points belonging to the neuron are all connected (Fig. 14).

There are many parameters in our reconstruction algorithm. The user should experiment with these parameters as the optimal settings may depend on properties of the images, which are likely to vary depending on staining and imaging procedures. Among the most important parameters are the distances between pixels and between the successive planes, which are determined by the image acquisition process. Also important is the number of subdivisions of one tiff stack. Since our algorithm relies on 2D projections, subdivision reduces overlap of neurites from different depths, and improves the reconstruction quality. Checking the validity of each SWC point is critical, so the users should pay close attention to adjusting these parameters. In Appendix 2, we have pointed out other important parameters while describing the technical details of the automatic reconstruction algorithm.

There are a number of open-source software packages for reconstructing neurons, most notably Vaa3D (Peng et al., 2014) and neuTube (Feng et al., 2015). Vaa3D has extensive capabilities for processing images from various kind of sources (Peng et al., 2014). In contrast, we focused on optimizing our software for the particular application of neurons stained with a dark reaction product following patch-clamp recording. Although ShuTu may work for neurons stained in other ways, we have made no attempt to optimize it for use with multiple staining and imaging procedures. In addition, ShuTu was developed with a philosophy that perfect automatic reconstruction is difficult, if not impossible. Therefore, we emphasized the importance of manual annotation and error correction. In keeping with this philosophy, ShuTu includes user-friendly software to facilitate these processes.

Another open-source software package for neuron reconstruction – neuTube – also has a strong 3D capability for manipulating SWC structure (Feng et al., 2015). As ShuTu is based on neuTube (T. Zhao is a contributor to both), many of the features of ShuTu are adaptations of neuTube. However, ShuTu includes several important extensions. neuTube was designed to deal with single tiff stack. As such, it was not designed to deal with reconstructing entire neuron, unless the neuron is contained in a single tiff stack.

ShuTu is a complete solution that includes the capability to deal with multiple tiff stacks, including modules for processing and stitching the images. In the interactive mode, the neuron structure is represented in multiples ways that are all linked (Fig. 11), thus improving the ease and accuracy of editing the SWC structure.

Commercial solutions for neuron reconstruction also exist (e.g. Neurolucida 360, MBF Bioscience; Imaris FilamentTracer, Bitplane). Detailed comparison of ShuTu to these other software packages is difficult, as it requires mastery of all of them to be fair. We encourage authors and users of other software packages to test ShuTu on their dataset or test their favorites on the images used in this work, and provide feedback.

Staining neurons with biocytin is common in patch-clamp experiments. However, methods for reconstructing neurons based on biocytin are limited. When dealing with bright-field images like the biocytin data, a common strategy is to apply some preprocessing method first (Türetken et al., 2011; Narayanaswamy et al., 2011; Zhou et al., 2015), making the images friendly for automatic reconstruction. Preprocessing, however, is often computationally intensive and does not guarantee good performance. ShuTu is specifically tailored to deal with inherent problems with images from biocytin filled neurons.

ShuTu is not restricted to biocytin-filled neurons. In principle, it can also handle images from confocal and fluorescent microscopy, simply by inverting the images. However, we made no attempt to develop this application of ShuTu. Care is likely to be necessary in ensuring that microscopy and image acquisition properties are optimized to maximize the utility of ShuTu for this application. For now, we have chosen to leave this enhancement to future iterations (by us or others) and focus our efforts on one common method of staining and imaging patch-clamped neurons.

Improving image quality will inevitably improve the efficiency and accuracy of neuron reconstruction. Users should ensure that high quality images are taken by following proper microscopy practices and protocols. Tissue fixation and clearing processes can influence the accuracy of the reconstructed neurons. Tissue shrinkage often occurs during the fixation process. To be accurate, these factors need to be quantified for specific experimental settings and the dimensions of the reconstructed neurons need to be adjusted to account for shrinkage and distortion.

Accurate estimates of diameters of dendrites are important for computational modeling of neurons (Psarrou et al., 2014; Anwar et al., 2014). Our automated algorithm works quite well in determining the diameters, as shown in Fig. 8b-c. Except for a few cases, manual adjustments of the diameters are rarely necessary during the manual editing stage. However, systematic biases can exist, especially if spines are densely stained and blurred in the images, for example in the Purkinje neuron shown in Fig. 16. Another common bias is due to tissue shrinkage during the fixation process. Users need to correct for these biases before using the SWC structures for simulations.

ShuTu has some limitations. It is not designed to trace axons, which are often too thin and faint following biocytin staining in slices to trace automatically. Spines are not marked. It is possible that editing operations currently requiring human judgements, such as when dendritic branches closely cross each other, could be automated in the future using machine learning approaches (Turaga et al., 2010).

In conclusion, we have shown that ShuTu provides a practical solution for efficient and accurate reconstructions of neuron morphology. The open-source nature of the software will allow the research community to improve the tool further, and increased efficiency in neuronal reconstruction should facilitate more studies incorporating quantitative metrics of dendritic morphology and computer simulations of dendritic function.

## Materials and Methods

### Whole-cell recording and neuron staining

All experiments were performed according to protocols approved by the Institutional Animal Care and Use Committee of the Janelia Research Campus.

Acute hippocampal slices were prepared from mice (17-30 days old). After animals were deeply anesthetized with isoflurane, they were decapitated and the brain rapidly removed into chilled cutting solution consisting of (in mM) 215 sucrose, 2.5 KCl, 20 glucose, 26 NaHCO_3_, 1.6 NaH_2_PO_4_, 1 CaCl_2_, 4 MgCl_2_, and 4 MgSO_4_. Hippocampi were dissected out and cut into 400 *µ*m thick transverse sections on a Leica VT 1200s vibrating microslicer (Leica, Ltd., Germany). The cutting solution was slowly exchanged with artificial cerebrospinal fluid (ACSF) containing (in mM) 124 NaCl, 2.5 KCl, 10 glucose, 26 NaHCO_3_, 1.0 NaH_2_PO_4_, 2.0 CaCl_2_, and 1.0 MgCl_2_. Both cutting and ACSF solutions were saturated with 95% O_2_ and 5% CO_2_ (pH 7.4). The slices were incubated at room temperature for at least 1 hour before recording, and then were transferred as needed to a submersion-type recording chamber perfused with ACSF at 2 ml/min.

Whole-cell recordings were obtained by visualized patch technique under IR-DIC optics. The recording pipette resistance ranged between 4 and 6 *M* Ω. Series resistance (6 − 15*M* Ω) and input resistance were monitored throughout each voltage-clamp recording. Recordings with >10% change in series resistance were excluded. All experiments were performed in the current-clamp configuration. The intracellular pipette solution consisted of (in mM) 135 K-gluconate, 5 KCl, 1 CaCl_2_, 0.1 EGTA-Na, 10 HEPES, 10 glucose, 5 MgATP, and 0.4 Na3GTP, 0.1% biocytin, pH 7.2 280-290 mOsm. Resting potential ranged from −69 to −58 mV. Maximal recording time after dissection was 6 hr. Recording temperature was set to 32.0 *±* 0.1 C^*◦*^ using a TC-344A single-channel temperature controller (Warner Instruments, Inc, Hamden, CT, USA). All experiments were executed with a Dagan BVC-700 amplifier, digitized (3 − 5 kHz) using an ITC-16 analog-to-digital converter (Instrutech) and analyzed using custom-made software for IgorPro (Wavemetrics Inc., Lake Oswego, OR, USA). All chemicals were purchased from Sigma-Aldrich (St. Louis, MO, USA). Neurons were filled with biocytin and fixed (12-24 hours) with paraformaldehyde (4%) after recording, then washed in 1X PBS solution. Biocytin staining was carried out with vector PK4000 and SK4100 kits (Vector Laboratories, Burlingame, CA, USA). Tiled z-stack images acquisitions was performed using a Zeiss AxioImager microscope with AxioCam and ZEN blue software at 100X or 63X magnification.

Sparse labeling of cerebellar Purkinje cells was achieved by in utero ventricular viral injection (100 nL per ventricle) at embryonic day 14 (E14) with an adeno-associated virus (Pseudo type 2.1) carrying a GFP payload expressed under the CAG promoter. Once animals reached postnatal day 30 (P30) they were transcardially perfused with 4% paraformaldehyde fixative. Fixed brains were then sectioned at 100 micrometer thickness (horizontal plane) using a Leica micro-slicer and mounted for microscopy. Tiled z-stack image acquisition was performed using a Zeiss 880 confocal microscope at 63X magnification.

### System requirements and installation

ShuTu consists of two parts: one for processing images and automated reconstruction, and the other for viewing and editing the morphology using graphical user interface (GUI). The software requires installation of Open MPI and C compiler. The software package was tested on a desktop computer with Intel Core i7-4770 CPU@3.40GHz CPU and 16 GB memory, running Ubuntu 14.04 LTS. These are typical settings for current high-end desktop computers. Multiple processors are desirable since the algorithms are designed to utilize multiple processors to speed up computation. However, the memory usage must be monitored to make sure that the demand on memory does not exceed 100%. The number of processors used is specified as a parameter for the command line when running the code for automated reconstruction (see the section “Automated reconstruction” below).

ShuTu can be downloaded from http://personal.psu.edu/dzj2/ShuTu/. or https://www.janelia.org/shutu,

An installation script is provided for Ubuntu and Mac OSX systems. This download includes all source codes for processing images and automatic reconstruction. It also includes the GUI program for viewing and editing the morphology. The source code for the GUI program is available at https://github.com/tingzhao/ShuTu.

On Windows 10, one can install the Ubuntu App and proceed as in Ubuntu, except that the GUI program is downloaded and installed separately and runs in Windows system.

In the directory of ShuTu, one can run

~~~
sudo sh build.sh,
~~~

which checks and installs necessary software including Open MPI. The C programs are also compiled.

### Image acquisition and processing

The software works with tiles of tiff stacks covering the entire neurons. Nearby tiles overlap, typically by 20%, to help fine tune the relative positions of the tiles (“stitching”). The names of the tiff stacks use the convention of a common string (`filenameCommon` followed by a number and. tif). With the *x, y* positions of the tiles specified, one can use the program `stitchTiles` to stitch the tiles. The results are stored in file `filenameCommon.json`.

Modern microscopes often allow automatic generations of overlapping tiles of tiff stacks. In our case, we imaged hippocampal neurons with Zeiss Axio Imager with AxioCam and Zen blue software. Once the boundary in the field of view (*XY*) and the range of the depths (*Z*) that contain the neuron are set, the images at each tile position and depths are automatically taken, and the positions of the image are stored in an xml file. The filenames of these images contain information about the tile number and depth. Using them, we assemble all images at different depths for each tile into one tiff stack. The command is

~~~
mpirun -n numProc. /createTiffStacksZeiss dirData filenameCommon
~~~

Here `numProc` is the number of processors to be used; and `dirData` is the path to the directory in which the `xml` file resides. The images of the planes are stored in a subdirectory. filenameCommon is the common part of the names given by the user to the created tiff stacks.

The user can generate the overlapping tiff stacks in other ways. The files should be named in the format of `filenameCommon1.tif, filenameCommon2.tif`, etc.

The tiff stacks are preprocessed using the command

~~~
mpirun -n numProc. /processImages dirData
~~~

If the images are dark-field, the command should be

~~~
mpirun -n numProc. /processImages dirData 1
~~~

In this case, the images are inverted into bright-field images. The original images are renamed by adding. `org.tif` to the end of the original file names, and are moved to a directory `OriginalImages`.

Stitching the images is done with program `stitchTiles`. It is assumed that in `dirData` there exists the `xml` file, generated by the Zen Blue software during automatic image acquisation; or a text file `tileSequences.txt`, with the following format explained in with an example:

~~~
80
1, 2, 3, 4
1, 5
5, 6, 7, 8
6, 9
9
~~~

The first line is a single number specifying the shifts in percentages (100 − percent overlap). The second line specifies the tile numbers of the first row scanning from left to right. The third line specifies two connected tiles in the first row and the second row. The fourth line specifies the tile numbers in the second row. This continues until the last row is specified. With one of these files in the directory, the command for stitching is

~~~
mpirun -n numProc. /stitchTiles dataDir
~~~

After stitching is done, one can proceed to reconstruct the neuron semi-automatically using the GUI program of ShuTu (see Appendix 1). Another choice is to run the automatic reconstruction algorithm to create a draft reconstruction and edit it using the GUI program (see below).

In stitching, the precise offsets of nearby tiles are computed by maximizing phase correlation (Zitova and Flusser, 2003). Using the maximum spanning tree algorithm (Graham and Hell, 1985), a tree graph connecting all tiles and maximizing the sum of phase correlations along the connected nearby tiles is computed and used to set the relative coordinates of all tiles.

If the entire neuron is contained in a single tiff stack, the above processing steps should be skipped.

### Automated reconstruction

The code for automated reconstruction is parallelized with MPI protocol, and runs with the command

~~~
mpirun -n numProc. /ShuTuAutoTrace dataDir ShuTu.Parameters.dat,
~~~

where `ShuTu.Parameters.dat` is a text file that contains the parameters. This creates an SWC file `filenameCommon.auto.swc in dataDir`, which can be loaded in `ShuTu` for manual editing (`File→Load SWC`).

To automatically reconstruct neurites in a single tiff stack, one can run

~~~
./ShuTuAutoTraceOneStack dataDir/filename.tif ShuTu.Parameters.dat.
~~~

The resulting SWC file is stored in `filename.tif.auto.swc in dataDir`. This should be used if the entire neuron is stored in a single tiff stack. It is also useful for tuning the parameters. Note that if the image is dark field, one needs to edit the parameter file and set the image type parameter to 1.

## Acknowledgements

Research for D.Z.J was supported by the Visiting Scientist Program at the Janelia Research Campus, Howard Hughes Medical Institute; and by the Huck Institute of Life Sciences at the Pennsylvania State University.

## Appendix 1: Editing commands for ShuTu

### Loading a project

A reconstruction project can be opened by clicking on `Open Project` icon or `File → Open Project`. In the directory of the neuron, there should be a file `filenameCommon.tiles.json`, which is created after stitching the tiff stacks. Clicking on it opens `Tile View`, in which the 2D projections of the tiff stacks are shown. The 2D projection of the neuron should be visible. If there is a previous reconstruction of the neuron, which is stored in a file `filenameCommon.swc`, it will be automatically loaded and overlaid onto the 2D projection. The SWC file generated by the automated algorithm, `filenameCommon.auto.swc`, can be loaded by selecting `File → Load SWC`.

Double clicking on any tile in the `Tile View` loads the corresponding tiff stack in `Stack View`. The loaded SWC points are overlaid onto the tiff stack. To go up and down in the *z*-dimension, use the right and left arrow keys. The functions of the arrow keys can be also performed with mouse wheel or track pad when available.

Clicking on `Make Projection` button creates 2D projection of the tiff stack. The user can specify the number of subdivisions used in the projection. All of the projections of the subdivisions are contained in the `Projection View`, which can be browsed with the left and right arrow keys.

The SWC structure is also displayed in `3D View`. It can be rotated with the arrow keys, and shifted with the arrow keys while pressing the `Shift` key.

In all views, zoom is controlled with + and - keys. After zooming in, different parts of the images can be navigated by pressing-dragging the mouse.

If the neuron is contained in a single tiff stack, load the tiff stack with `File → Open`. Other steps are the same as described above. Dark field tiff stack should be converted into bright field stack with `Tools → Invert Intensity`.

### Editing SWC points

The SWC structure can be edited in `Stack View, Projection View`, and `3D View`. All editing can be reversed by `Ctrl-z` (or `Command-z` in Mac). Colors of SWC points indicate their topological roles in the structure: yellow and blue indicate the end points of branches; green the branching points; and red the interior points. Lines between SWC points indicate their connectivity.

In `Stack View`, an SWC point is plotted with a circle at its *xyz* position in the tiff stack. The radius of the circle is the same as that of the SWC point. As the focus plane shifts away from the *z* of the SWC point, the circle shrinks with its color fading. This helps the user to visually locate the *z* of the SWC points and inspect whether the positions and radii of the SWC points match the underlying signals of the neurites in the tiff stack.

Extension is the most used editing function. In `Stack View`, it can be done in two ways. The first is manual extension. Click an SWC point to extend, and the cursor becomes a circle connected to the SWC point. Focus on the target neurite using the arrow keys, and match the radius of circle with that of the neurite using e and `q. Ctrl`-clicking on the target points creates a new SWC point connected to the starting SWC point. (In Mac, use `Command` instead of `Ctrl`.) The second is smart extension. It is the same as manual extension, except that the user clicks without pressing `Ctrl`. This method allows clicking far from the starting SWC points; the algorithm fills in additional connected SWC points along the neurites with the radii and depths automatically calculated. Smart extension works well when the underlying signal is strong.

To change the properties of a particular SWC point, select it by clicking on it and pressing `Esc` to come out of the extension mode. The radius can be changed with `e` and `q`. It can be moved with `w,s,a,d` for up, down, left, right. Pressing x deletes it.

To connect two SWC points, click on the first point and `Shift`-click on the second point, then press c. Pressing `Shift`-c after selecting two points automatically fills additional SWC points, similarly as in the smart extension. To disconnect two SWC points, select them then press `b`.

In `Projection View`, 2D projections of the subdivisions of the tiff stack are overlaid with the SWC points. In this view it is easier to spot missed branches and incorrect connections. There is also a mask-to-SWC method for tracing branches. To draw a mask along a branch, press `r`. The cursor becomes a red dot. Roughly match the radius of the dot with that of neurite with `e` and `q`. Click on the start point, then `Shift`-click on the target. A red mask will be drawn along the branch. Clicking on `Mask → SWC` button converts the mask into SWC points, which can be examined in detail in the `Stack View`. The mask can also be drawn manually by press-dragging the mouse along the branch. To get of out the mask drawing mode, press `Esc`.

Clicking on an SWC point selects it. Pressing `z` locates the selected point in the `Stack View`, and its z position and other properties can be further examined with the tiff stack.

The user can directly modify the connections in the `Projection View`. The operations are the same as in the `Stack View`.

In `3D View`, the user can examine and modify the connections between SWC points. Connecting or breaking connections between two SWC points is the same as in the `Stack View` and the `Projection View`. Selecting an SWC point and pressing `z` locates it in the `Stack View` for further examination and extension. This operation also loads a new tiff stack if the selected point is not in the current tiff stack.

A useful way of locating broken points in the SWC structure is the operation that selects all connected SWC points to the selected SWC point. It is done by pressing `h-3`, or right-clicking the mouse and selecting `Select → All connected nodes`.

After correctly connecting all SWC points belonging to the neuron, the user can delete all noise points simply by selecting all SWC points in the neuron, right-clicking the mouse, and performing `delete unselected`.

### Annotating, saving, and scaling the SWC structure

After the reconstruction is done, the user needs to annotate the SWC points as soma, axon, apical dendrite, basal dendrite. This is best done in `3D View`. In the panel `control and settings`, change `Color Mode` to `Branch Type` to reveal the types of SWC points. To annotate the soma, the user can select one point in the soma, right-click the mouse, and select `Change Property → Set as root`. More SWC points belong to the soma can be selected by `Shift`-clicking. Then right-click to bring up the menu, then select `Change type` and set the value to 1. The SWC points in the soma are shown in blue.

To annotate the axon, select the one SWC point closest to the soma, and press `h-1`. This selects all SWC points down stream of the selected point. Then change type to 2. Basal dendrites and apical dendrite can be similarly annotated, and their types are 3 and 4, respectively.

In the panel `control and settingings`, setting `Geometry` to `normal` produces the volume representation of the SWC structure, with surface rendered between adjacent SWC points.

To save the reconstruction, click on the objects in the panel `Objects`, which selects the corresponding SWC points. Then in the window of the SWC structure, left-click and do save as. It is best to use the default filename `filenameCommon.swc`.

The dimensions of the SWC points in `filenameCommon.swc` are pixel based. To convert them into physical dimensions in *µ*m, type in the terminal

~~~
./scaleSWC dataDir ShuTu.Parameters.dat
~~~

This process uses `xyDist` and `zDist` in `ShuTu.Parameters.dat`, which specify in *µ*m the *xy* pixel distance and *z* distance between successive planes. The results are saved in `filenameCommon.scaled.swc`.

Right after finishing the reconstruction and with ShuTu closed, the number of various editing operations can be analyzed using the command

~~~
python analyzeNEO.py
~~~

This requires the user to install Python 2.7. A plot similar to Fig. 15c will be generated. The script `analyzeNEO.py` parses the log file generated by `ShuTu`. The log file can contain several neuron reconstruction sessions, but the script only parses the most recent one. When estimating the total time of manual editing, idle times of the user are excluded if they are detected in the log file. The log file is assumed to be at

~~~
∼/.neutube.z/log.txt
~~~

If the log file is in other locations, the user can use command

~~~
python analyzeNEO.py logFileDir/log.txt.
~~~

There are many more editing functions in ShuTu. The user can refer to Help for more instructions.

## Appendix 2: Technical details of automated reconstruction

Here we provide technical details of the automated reconstruction algorithm presented in the main text. These details should help the users to adjust parameters for their specific needs, and facilitate further development of the algorithm. The parameters in each step are summarized in series of tables. The algorithm is explained with the same example used in the main text.

### Coordinate system

A tiff stack consists of successive 2D images (referred to as planes) taken at increasing depths at regular intervals. We denote a pixel in a tiff stack with coordinates (*x, y, z*). Here *x, y* are the pixel positions in the planes, and *z* is the depth. We take the convention that in a plane, the *x* axis points vertically downwards and the *y* axis horizontally to the right (Fig. 1).

The distance between neighboring pixels in *x* and *y* is denoted as *d*_*xy*_. The distance between successive planes is denoted as *d*_*z*_. In the example, *d*_*xy*_ = 0.065 *µ*m (`xyDist`, name in the parameter file), and *d*_*z*_ = 0.5 *µ*m (`zDist`; Table 1).

**Table 1:** Parameters for tiff stacks and 2D projections. Names of the parameters appear in the file containing parameters for automated reconstruction.

| Parameter | Name | Value | Meaning |
| --- | --- | --- | --- |
| $d_{xy}$ | **xyDist** | $0.065 \mu\text{m}$ | pixel distance in xy |
| $d_z$ | **zDist** | $0.5 \mu\text{m}$ | z-distance between successive planes |
| $\sigma_b$ | **sigmaBack** | $2 \mu\text{m}$ | length scale for smooth background |

### Prepocessing

Our algorithm requires that the images are grayscale with bright background. Other image types must be converted into bright-field grayscale images, and this is done in preprocessing. In particular, color images are converted into grayscale according to

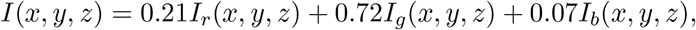

where *I* is the intensity of the grayscale and *I*_*r*_, *I*_*g*_, *I*_*b*_ are those of the red, green, and blue channels. Dark-field images are inverted by subtracting the grayscale intensity at each pixel from the maximum intensity of the tiff stack. To reduce pixel noise, each plane is smoothed with 2D Gaussian filter with *σ* = 1 pixel. The intensity is scaled so that the maximum is 1 for the tiff stack.

### 2D projection

We identify neurites in a tiff stack from its minimum-intensity 2D projection. The intensity *I*(*x, y*) of the 2D projection is taken as the minimum intensity among all pixels with the same *z*. Projections of dendritic branches form dark paths in *I*(*x, y*) (Fig. 3a). Shadows of branches in out-of-focus planes (Fig. 2d) do not create separate dark paths in the 2D projection; instead, their projections flank those of the branches, forming smooth decay of intensity away from the center lines of the branches. The problem of confusing the shadows of the branches as neurites in the out-of-focus planes, as shown in Fig. 2d, does not exist in the 2D projection.

To eliminate smooth variations of the background due to uneven lighting, we subtract from *I*(*x, y*) a background, which is obtained by blurring *I*(*x, y*) with a Gaussian filter with standard deviation *σ*_*b*_ = 2 *µ*m (sigmaBack; Fig. 3b). We then normalize the range of *I*(*x, y*) to (0, 1) (Fig. 3c). Smaller *σ*_*b*_ enhances weak signals relative to strong signals (Fig. 3d). This is because the background with smaller *σ*_*b*_ tracks the signal strength more closely, and when subtracted, takes away more from the strong signals. But *σ*_*b*_ should be large enough to ensure that the subtracted background is smooth and does not weaken the signals.

### Binary mask

From the 2D projection we create a binary image *b*(*x, y*) to indicate pixels that belong to neurites. Specifically, *b*(*x, y*) = 1 for pixels in the neurites (foreground pixels) and *b*(*x, y*) = 0 for those in the background (background pixels). We call the area defined by the foreground pixels as binary mask.

The first step in creating the mask is convolving *I*(*x, y*) with valley detectors with varying orientations, and finding the maximum and minimum responses to the detectors (Fig. 3e). A valley detector *f* (*x, y*) is a patch of 2D image (or filter) consisting of an oriented dark band flanked by two bright bands. Mathematically the filter is expressed as

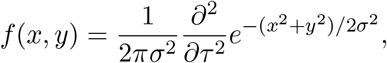

which is a directional second derivative of a Gaussian with standard deviation *σ*. Here

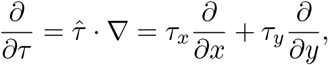

where 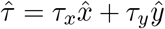 is a unit vector perpendicular to the orientation of the dark band.

Convolving *I*(*x, y*) with the filter creates the response *R*(*x, y*):

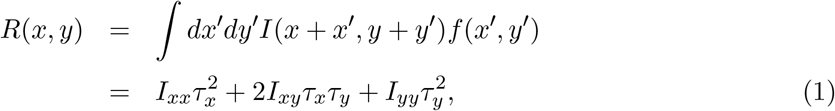

where

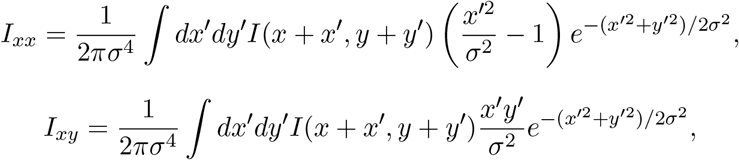

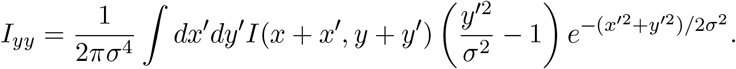

We obtain the maximum or minimum response at (*x, y*) using the Lagrange multiplier method:

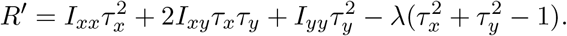

At the extrema we have

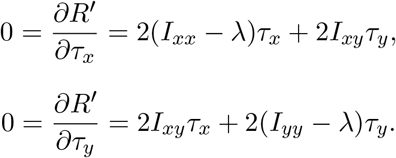

These are linear equations, which can be expressed in matrix form as

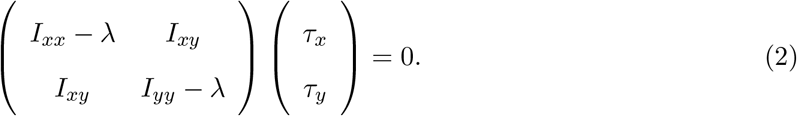

To have none-zero solutions for *τ*_*x*_ and *τ*_*y*_, we must have

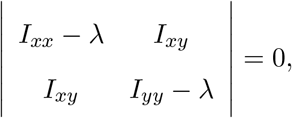

where *λ* is the eigenvalue of the Hessian matrix. There are two solutions:

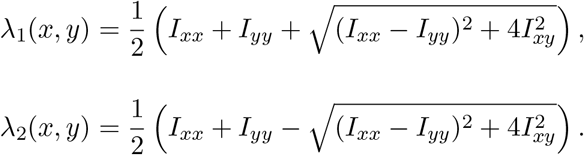

Here we chose *λ*_1_(*x, y*) > *λ*_2_(*x, y*). Solving *τ*_*x*_ and *τ*_*y*_ from Eq. (2) and plugging in to Eq. (1), we find the response at the extrema:

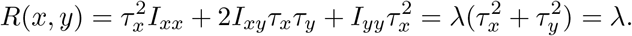

Hence the maximum *R*_*m*_(*x, y*) of the responses *R*(*x, y*) to valley detectors at varying orientations is given by

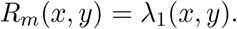

To see how we can create the mask from *λ*_1_(*x, y*) and *λ*_2_(*x, y*), we exam three simple examples of synthetic 2D images containing some aspects of 2D projections of the real images containing neurites.

The first example is a Gaussian valley in *y* direction:

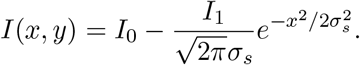

Here *σ*_*s*_ is the scale of the widths of the valley; *I*_0_ is the baseline intensity; and *I*_1_ is the amplitude. This is an idealized model of the 2D projection of a dendritic segment with half-width *σ*_*s*_. An ideal mask for this Gaussian valley is a rectangular strip spanning the *y* direction, centered long *y*-axis and with half-width *σ*_*s*_.

*λ*_1_(*x, y*) and *λ*_2_(*x, y*) are easily calculated. We find that

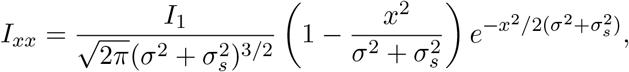

and

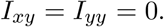

Therefore,

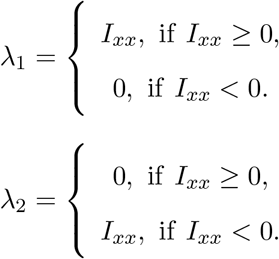

We can obtain a mask close to the ideal mask by thresholding *λ*_1_(*x, y*). If we set the foreground pixels as those with *λ*_1_(*x, y*) > 0, the boundary of the mask is given by

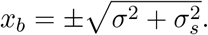

The half-width of the mask is 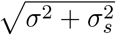, and it is larger than *σ*_*s*_. Taking *σ →* 0 leads to the ideal mask. For finite *σ*, it is possible to set a higher threshold for *λ*_1_(*x, y*) and obtain the ideal mask; but this requires a threshold that depends on the width of the valley.

From this example we see that we can obtain a mask that closely follow dendritic branches by thresholding the maximum responses to the valley detectors, *λ*_1_(*x, y*). The threshold should be larger than 0. Larger *σ* for the detectors tends to broaden the mask; therefore it is desirable to have small *σ* to obtain masks that closely cover the dendritic branches.

The second example is a Gaussian blob:

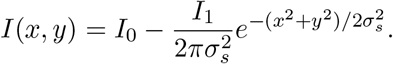

This is an idealized model for the 2D projections of spills created during the staining process (Fig. 2a). Such spills are noise that should be eliminated; therefore the ideal mask for a Gaussian blob should be empty.

We find that

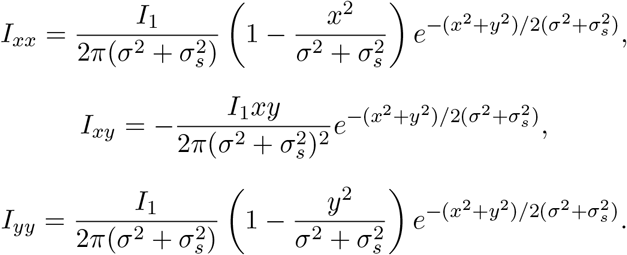

Therefore,

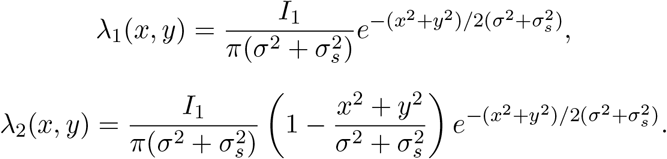

We see that thresholding the maximum responses *λ*_1_ creates a circular mask, which is far from the desired empty mask. To suppress creating foreground pixels for the Gaussian blob, additional criteria for the mask are needed. We notice that near the center of Gaussian blob, *λ*_1_ and *λ*_2_ are approximately equal. This motivates another criterion for the mask: in addition to *λ*_1_ being greater than a threshold, the foreground pixels must satisfy the condition *λ*_1_ > *α_λ_|λ*_2_*|*, where *α_λ_* > 1 is a factor. This criterion should suppress foreground pixels for the Gaussian blob, except around a ring near the radius 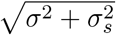, where *λ*_2_ is close to zero. This is not the ideal mask for Gaussian blob, but it is close. Note that this additional criterion does not affect the mask for the Gaussian valley in the first example, hence does not interfere with detection of dendritic branches.

The third example is a random image, which has a mean intensity *I*_0_ and no correlations between the pixels:

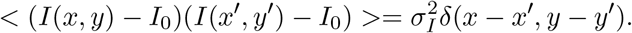

Here 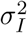 is the variance of the pixel intensity. This is an idealized model for random pixel noise in the real images. The ideal mask should be empty.

It is easy to see that

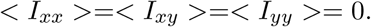

Additionally,

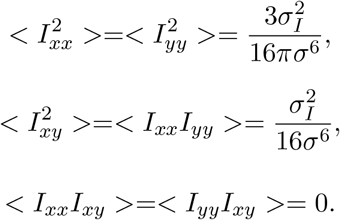

From these we find the mean of the responses

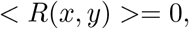

and the variance

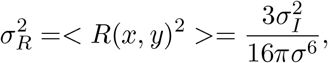

where we have used 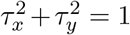. *σ_R_* represents the range of the responses expected from random fluctuations of the intensity. To avoid creating foreground pixels for random fluctuations, we should set the threshold for *λ*_1_ larger than *σ_R_*. But a threshold that is too large diminishes the mask for dendritic branches. Therefore the threshold for *λ*_1_ must be chosen to preserve the signals while suppressing random noise. Inevitably, some foreground pixels to noise are unavoidable, which creates random speckles in the mask (Fig. 3h).

A guideline for selecting the length scale *σ* in the valley detectors can be devised by combining the insights from the Gaussian valley and the random image. The peak of the maximum responses from the Gaussian valley, which occurs at *x* = 0, is given by

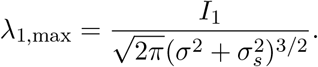

Comparing this to the variance of the responses to the random image, we can define the signal-to-noise ratio as

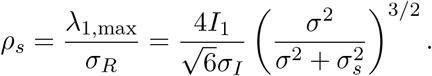

This ratio is an increasing function of *σ*. Therefore, a large *σ* is useful for suppressing noise. However, a large *σ* overestimates the width of the valley (given by 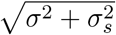), leading to widening of the foreground pixels. For real images, such widening can create a mask that merges nearby branches, leading to an inaccurate representation of the neuronal structure. Hence the choice of *σ* is a compromise between enhancing the signal-to-noise ratio while avoiding the merger of nearby branches in the mask. When the intensity fluctuation is small, we can select a small *σ*, leading to an accurate mask. If the fluctuation is large, we need to choose a large *σ* and live with the imperfect mask.

Based on the insights gained from the examples discussed above, we formulate the following procedure for creating the mask *b*(*x, y*) from *λ*_1_(*x, y*) and *λ*_2_(*x, y*). Select *σ* of the valley detector such that the neurites are clearly visible in *λ*_1_(*x, y*) (Fig. 3d). Set *b*(*x, y*) = 0 with *λ*_1_ < *α_λ_|λ*_2_*|* to suppress circular blobs in the 2D projection. Select a threshold *θ_λ_* above the noise level, and set *b*(*x, y*) = 1 if *λ*_1_(*x, y*) > *θ_λ_* and *b*(*x, y*) = 0 otherwise (Fig. 3e). Since pixels belonging to the neurites tyically have higher *λ*_1_ compared to those with random fluctuations, we set the threshold *θ_λ_* such that the fraction of pixels selected to the mask is *f_λ_*. For our example neuron, the parameter values are: *σ* = 0.1 *µ*m (`sigmaFilter`), *α_λ_* = 10 (`lambdaRatioThr`), and *f_λ_* = 0.1 (`sparse`; Table 2).

**Table 2:** Parameters for creating the binary mask.

| Parameter | Name | Value | Meaning |
| --- | --- | --- | --- |
| $\sigma$ | **sigmaFilter** | $0.1 \mu\text{m}$ | length scale of valley detector |
| $\alpha_\lambda$ | **lambdaRatioThr** | 10 | factor for eliminating circular blobs |
| $f_\lambda$ | **sparse** | 0.1 | fraction for determining the threshold |
| $\mu$ | **levelSetMu** | 0.1 | parameter for level set smoothing |
| $N_{levelset}$ | **levelSetIter** | 500 | number of level set iterations |
| $A_s$ | **smallArea** | $1.0 \mu\text{m}^2$ | threshold for small areas removed |

The binary mask generated as above is noisy, and the boundaries for neurites are rugged (Fig. 3h). To clean up noise and smooth the boundaries, we use the sparse-field level-set method outlined in (Lankton, 2009), which is a technical report based on (Whitaker, 1998). The details of level-set smoothing is as follows.

For the 2D projection *I*(*x, y*) after background subtraction and normalization, we compute the gradient

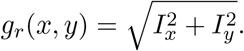

We rescale the gradient so that the range is from 0 to 1. An edge indicator is defined as

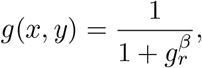

where *β* is an exponential, typically smaller than 1, for compressing the gradient values. This function is minimal at edges of branches, where the gradients are larger. We seek a contour *C* such that the energy function

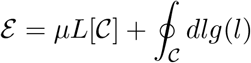

is minimized, where *L*[*C*] is the total length of the contour and *µ* is weight parameter that controls the smoothness of the contour. The curve that minimize this energy function will be smooth and sit along the maximum gradient boundaries between the branches and the background.

The contour can be expressed as the zero-crossing points of a level set function *ϕ*(*x, y*). Inside 𝓒, we have *ϕ* > 0, and outside *ϕ* < 0. Note that

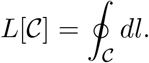

The unit vectors normal to the contours in *ϕ* are given by

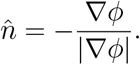

Hence

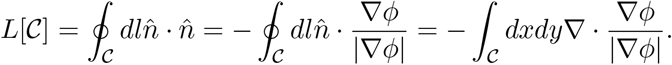

The last step uses the divergence theorem. Note that

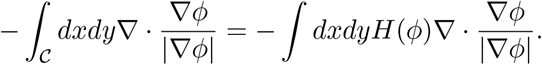

Here *H*(*ϕ*) is the step function; it is 1 if *ϕ* > 0 and 0 if *ϕ* < 0. Integration by part gives

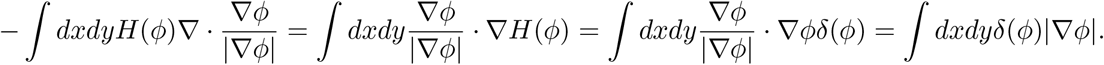

The surface term is zero because *H* is zero at the boundary. Here *δ*(*ϕ*) is the Dirac *δ*-function. Therefore, we have

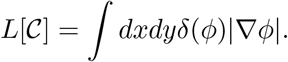

Similarly, we can derive

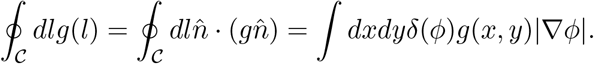

Hence, we have

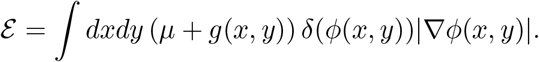

We use the variational method to find the *ϕ* that minimizes *ε*. Noting that

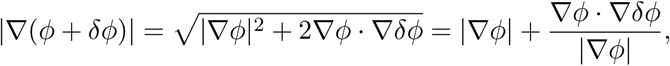

we find

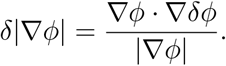

Applying integration by part, we have

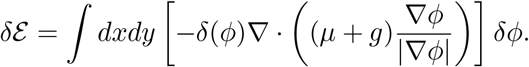

Setting *ε* = 0, we find

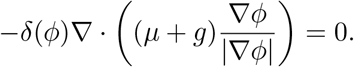

The surface term in the integration vanishes if we impose the Neumann boundary condition

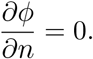

At equilibrium and on *𝓒* we have

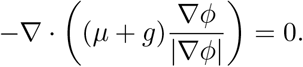

At other points, *ϕ* can be arbitrary. Minimization of

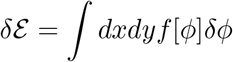

can be done by solving the equation

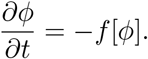

This equation implies

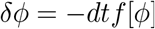

at each time step. Therefore

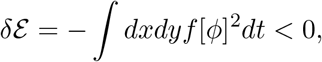

leading to decreasing *ε*. In our case, we need to evolve

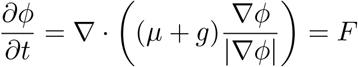

on the boundary. For *ϕ* at points other than the boundary, we need to change *ϕ* such that it remains a smooth function around the boundary and the second derivatives can be computed. We used the sparse-field implementation for solving this differential equation, which iteratively updates the sets of points near the boundary (Lankton, 2009). After *N*_*levelset*_ number of iterations, we obtain a new binary mask by setting *b*(*x, y*) = 1 if *ϕ*(*x, y*) > 0, and *b*(*x, y*) = 0 otherwise.

In practice, we observe that it is sufficient to smooth the initial mask by minimizing the length of the boundary alone. Hence we set the edge indicator

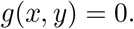

This is because the boundaries of the initial mask are already near the neurite boundaries.

The parameter *µ* (`levelSetMu`) controls the smoothness of the boundary. Smoothing deletes small noisy speckles. Larger *µ* creates smoother boundaries but can also cause small neurites to disappear. We set *µ* = 0.1. Also important is the number of iterations *N*_*levelset*_ (`levelSetIter`). It should be large enough to reduce noise and smooth the boundaries, but small enough not to loose structures due to over-smoothing. In our example we set *N*_*levelset*_ = 500.

As the final step, we remove connected pixels with total area smaller than *A*_*s*_ (`smallArea`). This removes noise and cleans up the mask (Fig. 3f). In the example we set *A*_*s*_ = 1 *µ*m^2^.

Parameters for creating the mask are listed in Table 2.

### Creating SWC points from the mask

Using the mask, we create the SWC points that describe the dendritic structure. The (*x, y*) positions of the SWC points are placed along the centerlines of the mask. The radii *r* are set as the shortest distances to the boundaries of the branches from the centerlines. The *z* positions are computed using the centerlines and the tiff stack.

The centerlines of the mask are obtained by skeletonization (Zhang and Suen, 1984). The skeleton is computed by iterative thinning of the mask based on the pixel values in the 8 neighboring points (Zhang and Suen, 1984) (Fig. 4a). The distance from a pixel in the centerline to the nearest boundary is computed using the Euclidian distance transformation (Danielsson, 1980) (Fig. 4b).

The depths *z* of the points on the centerlines of the mask are found in two steps. First, the centerlines are dissected into *xy*-paths (Fig. 4a). The *xy*-paths with length smaller than *l*_*sm*_ = 0.5 *µ*m (smallLen) are considered as noise and excluded. Second, a *z*-image is created by following the *xy*-path and cutting through the tiff stack (Fig. 4c). A dark valley in the *z*-image spanning from the left edge to the right edge indicates the branch whose 2D projection falls on the *xy*-path (Fig. 4c). A line through the valley can be found by evaluating all paths from the left edge to the right edge (red dotted line in Fig. 4c). Specifically, a left-right path in the *z*-image starts from a point at the left edge. The next point is selected from the three nearby points to the right (the change in *z* is −1, 0, or 1). This process iterates until the right edge is reached. For each left-right path, we compute the weighted distance, which is the sum of the distance between consecutive pixels multiplied by a weight *e*^*αdI*^, where *α_d_* = 20 (alphaDistance) is a parameter and *I* is the intensity of the right point. The weight penalizes bright pixels, and encourages the left-right path to go through dark pixels. The left-right path with minimal weighted distance, or the shortest path, is selected. The path follows the dark valley spanning from the left edge to the right edge, as shown in Fig. 4c (dotted red line). The *z* values of this shortest path gives the depth for each point in the *xy*-path. A large *α_d_* ensures that the shortest path follows dark pixels. But a value too large can lead to distortions of the path due to dark spots from other branches or noise.

Linked SWC points are placed on the *xy*-path. The distance between successive SWC points is set to be at least 1.2 times the radii (for points near som, 0.5 times radii). We check the validity of SWC points, remove any invalid ones. Adjacent SWC points are connected along the *xy*-path. Removal of invalid SWC points may have created a large distance between two consecutive SWC points; in this case, we do not connect them. The criterion is

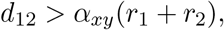

where *d*_12_ is the Euclidian distance in *xy* between the two SWC points; *r*_1_, *r*_2_ are the radii; and *α*_*xy*_ = 2.0 (`distFactConn`) is a factor. Sharp turns in *xy*-path often result from errors in the skeleton due to crossing branches. To avoid this problem, we do not connect the two SWC points if doing so creates a lage angle (greater than *θ_thr_* = *π/*3, `angle`) between consecutive lines connecting the SWC points. We also do not connection the SWC points if the *z* difference between them is too large, as this often leads to errors in connecting branches far way in *z*. The criterion is

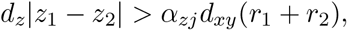

where *d*_*z*_ = 0.5 *µm* is the distance between successive planes in the stiff stack; *d*_*xy*_ = 0.065 *µm* is the pixel distance in *xy*; and *α_zj_* = 3.0 (zJumpFact) is a factor for adjusting the threshold.

The parameters for creating SWC points are listed in Table 3.

**Table 3:** Parameters for creating SWC points from 2D mask.

| Parameter | Name | Value | Meaning |
| --- | --- | --- | --- |
| $l_{sm}$ | **smallLen** | $0.5 \mu\text{m}$ | smallest length of xy-path used |
| $\alpha_d$ | **alphaDistance** | 20 | factor for weighting distance between pixels |
| $\alpha_{xy}$ | **distFactConn** | 2 | factor for disconnecting two consecutive SWC points |
| $\theta_{thr}$ | **angle** | $\pi/3$ | maximum angle allowed between consecutive lines |
| $\alpha_{zj}$ | **zJumpFact** | 3 | factor for z jump threshold |

### Checking the validity of an SWC point

The SWC points created from the mask can be incorrect. For example, if some branches are parallel to each other and are very close, they can be merged in the mask, leading to incorrect SWC points. Therefore it is important to check the validity of the SWC points and reject incorrect ones. Since the mask can be imperfect, we check the validity not with the mask but with the original tiff stack. The main idea is that a valid SWC point should sit on the centerline of a valley in the plane at the depth of the SWC point.

Consider an SWC point at (*x*_*p*_, *y*_*p*_, *z*_*p*_) with radius *r*_*p*_. The validity of the point is tested with the intensity *I*(*x, y*) of pixels in the plane at *z* = *z*_*p*_ (Fig. 5a). We take a local square patch in the plane centered at (*x*_*p*_, *y*_*p*_) with size set to max(4*r*_*p*_, 2*r*_*min*_), where *r*_*min*_ = 2 *µ*m (minRange). Pixel intensity across the patch can have a tilt, such that one side is much brighter than the other. This could be due to uneven lighting, or shadows cast by nearby dark neurites. To reduce the adverse effects of tilt, we least-square fit the intensity with a linear function *a*(*x* − *x*_*p*_) + *b*(*y* − *y*_*p*_) + *c*, where *a, b, c* are the fitted parameters, and subtract the fitted function from *I*(*x, y*) in the patch. The results are shifted and scaled so that the intensity range is the same as that before subtraction.

Ideally, the SWC point should be at a local center of a valley in *I*(*x, y*) of a dendritic branch. To test this, we create a profile of intensity along a line through (*x*_*p*_, *y*_*p*_) and at angle *θ* relative to the *x*-axis (Fig. 5a), and test the existence of an inverse peak. The profile is a one-dimensional curve *I*_*θ*_(*d*) (black line, Fig. 5b), where *d* is the coordinate of a pixel point on the line, with (*x*_*p*_, *y*_*p*_) set as the origin. *I*_*θ*_(*d*) is the intensity value at the pixel point. The range of *|d|* is limited to *d*_*max*_ = min(2*r*_*p*_, *r*_*min*_). We obtain a smoothed profile *I*_*s,θ*_(*d*) by convolving *I*_*θ*_(*d*) with a Gaussian filter with *σ*_*s*_ = 0.2 *µm* (`sigmaSmoothCurve`, green line, Fig. 5b), and detect the inverse peak in *I_s,θ_*. We take *θ* to be multiples of *π/*8. Hence there are 8 profiles (Fig. 5a,c).

We evaluate the existence of an inverse peak in the smoothed profile *I*_*s,θ*_ using two criteria. The first criterion ensures that the inverse peak is deep enough. Specifically, we require that the local minimum of *I*_*s,θ*_ near (*x*_*p*_, *y*_*p*_) is smaller than a threshold *I*_*th*_ = *I_b_ − α_th_σ* (gray line, Fig. 5b). Here *I*_*b*_ is the baseline, and is set to 80 percentile value of the intensity in the patch; *σ* = 0.03 (`sigma`) is the estimate of the fluctuation level of the intensity; and *α*_*th*_ = 1 (`factSigmaThreshold`) is an adjustable factor. This factor is increased to *α*_*th*_ = 2 (`factSigmaThreStrict`) during creation of SWC points from the mask to make the validity judgement stricter, which reduces noise in the SWC structure. The second criterion ensures that the inverse peak has two flanks. This is done by checking that *I*_*s,θ*_ rises above a threshold set at the mean of the maximum and the minimum of *I*_*s,θ*_ at either side of the local minimum point (gray dashed line, Fig. 5b).

If the peak is deep enough and there are two flanks, we determine the width of the peak starting from the minimum of *I_s,θ_*. We take a derivative of *I_s,θ_* and smooth it to get *dI_s_* (Fig. 5e). We determine the maximum absolute value of these derivatives *dI*_max_. Starting from the minimum point, we trace the negative part of the derivative. The tracing stops once *dI_s_* starts to increase and *−dI_s_* > 0.5*dI*_max_. This way of stopping ensures that the tracing picks out the first significantly large absolute value in the derivative and does not stop because of small bumps in *dI*_*s,θ*_. Similarly, we trace the positive part of the derivative, and stops when *dI_s_* starts to decrease and *dI*_*s*_ > 0.5*dI*_max_. If either of these tracing does not stop before reaching the end, the peak is invalid. Otherwise, the width *w* of the peak is set to the distance between the two stopping points (black vertical lines, Fig. 5e). The derivatives of all eight smoothed profiles are shown in Fig. 5f. To ensure that fluctuations between the two stopping points are small enough, we check the peaks in *|dI_s_|*. If the maximum of these peaks is larger than 0.5*dI*_max_, the profile is judged to have no valid peak.

If none of the profiles have valid inverse peaks, the SWC point is invalid. Otherwise, we chose the profile with the minimum width among the valid ones, and assign its peak position as the position (*x*_*m*_, *y*_*m*_) of the SWC point, and set *r*_*m*_ = *α*_*r*_*w/*2, where the factor *α*_*r*_ = 1.0 (`factAdjustRadius`) is an adjustable factor introduced to enable the user to adjust the radii of SWC points according to subjective judgement. If the distance between the original position (*x*_*p*_, *y*_*p*_) and (*x*_*m*_, *y*_*m*_) exceeds *α*_*shift*_*r*_*m*_, where *α_shift_* = 2 (`factShift`), then the SWC point has shifted too much, and it is flagged as invalid. If *r*_*m*_ is smaller than *r*_*min*_ = 0.3 *µ*m (`minRadius`) or larger than *r*_*max*_ = 10 *µ*m (`maxRadius`), the radius of the SWC point is too small or too large, and the point is invalid.

Finally, we check whether the pixels within a radius 0.5*r*_*m*_ from the center are dark enough. Specifically, we check whether the maximum value of the smoothed profile in the orthogonal direction (red line,Fig. 5d) to the chosen profile (cyan line,Fig. 5d) within 0.5*r*_*m*_ are smaller than a threshold (green horizontal line, Fig. 5d), which is set as the maximum smoothed intensity of the chosen profile at *r*_*m*_ plus *σ*. If not, the SWC point in invalid. This check ensures that the SWC points created along edges of thick dendrites or the soma are eliminated.

If the SWC point passes all the tests described above, it is judged as valid, and (*x*_m_, *y*_*m*_, *z*) and *r*_*m*_ are set as the new position and radius. The position and radius of the SWC are thus corrected after the validity check. To ensure that the corrections converge, the checking procedure is iterated three times with the positions and radius updated. The SWC point is accepted if it passes the tests all three times. The angle *θ* of the profile denotes the orientation perpendicular to the branch (red line, Fig. 5a).

The parameters used in checking the validity of an SWC point are listed in Table 4.

**Table 4:** Parameters for checking validity of an SWC point.

| Parameter | Name | Value | Meaning |
| --- | --- | --- | --- |
| $r_{min}$ | **minRange** | 2 $\mu\text{m}$ | lower bound for the range of the profiles |
| $\sigma_s$ | **sigmaSmoothCurve** | 0.2 $\mu\text{m}$ | length scale for smoothing profiles |
| $\sigma$ | **simga** | 0.03 | estimate of fluctuations in the intensity |
| $\alpha_{th}$ | **factSigmaThreshold** | 1 | factor for determining the threshold for peak |
| $\alpha_{th,s}$ | **factSigmaThresholdStrict** | 2 | strict factor for the threshold for peak |
| $\alpha_{shift}$ | **factShift** | 2 | factor for allowing shifts in the position |
| $r_{min}$ | **minRadius** | 0.2 $\mu\text{m}$ | lower bound for the radius |
| $r_{max}$ | **maxRadius** | 10 $\mu\text{m}$ | upper bound for the radius |
| $\alpha_r$ | **factAdjustRadius** | 1.0 | factor for adjusting the radius |

### Adjusting *z*

Checking validity of an SWC point shifts its *xy* position. The shift can be significant, especially during the creation of SWC points using the mask, since the mask can deviate from the underlying neurites significantly if there are close branches. To ensure that *z* of the SWC point is accurate, we further adjust *z*. We create the intensity profile along *z* at the *xy* position (black line, Fig. 4c), then smooth it with parameter *σ*_*s,z*_ = 2 (`sigmaSmoothCurveZ`; green line, Fig. 4c).

We check the existence of inverse peak in the smoothed profile, similarly as done for the intensity profiles used for checking validity of SWC points. We estimate the fluctuations in the profile by computing the standard deviation *σ_z_* of the difference between the original and the smoothed profiles. We smooth the derivative of the smoothed profile, and determine the maximum *d_z,max_* of the absolute values of the derivatives. If *d*_*z,max*_ < *α*_*z*_*σ*_*z*_, it is judged that there is no inverse peak, and the SWC is judged as invalid. Here *α*_*z*_ = 0.1 (`factSmallDerivZ`) is a factor. Otherwise, the extent of the inverse peak centered around the original *z* position is decided by finding the first places away from the center at which the smoothed profile stops increasing. A threshold is set to the maximum value at these two ends minus *α*_*th,z*_*σ*_*z*_. Here *α*_*th,z*_ = 1 (`factSigmaThresholdZ`) is a factor. If the minimum value within the extent is smaller than this threshold, the inverse peak is valid, and the *z* is adjusted to the position of the minimum. Otherwise the SWC is judged invalid.

The parameters for adjusting *z* are listed in table 5.

**Table 5:** Parameters for adjusting *z* of an SWC point.

| Parameter | Name | Value | Meaning |
| --- | --- | --- | --- |
| $\sigma_{s,z}$ | **sigmaSmoothCurveZ** | 2 | parameter for smoothing $z$ profile |
| $\alpha_z$ | **factSmallDerivZ** | 0.1 | factor for determining the significance of derivatives |
| $\alpha_{th,z}$ | **factSigmaThresholdZ** | 1 | factor for determining the threshold for inverse peak |

### Mark pixels occupied

To avoid creating duplicated SWC points, we mark pixels in the tiff stack in the vicinity of the existing SWC points as occupied. Before creating a new SWC point, we check whether the pixel at its center is marked as occupied; if so, the SWC point is not created.

The marked pixels around two connected SWC points (*x*_1_, *y*_1_, *z*_1_, *r*_1_) and (*x*_2_, *y*_2_, *z*_2_, *r*_2_) are in a volume formed by two half cylinders with radii *α_occ_r*_1_ and *α_occ_r*_2_, respectively, and a trapezoidal prism that fits with the half cylinders (Fig. 6). Here *α_occ_* = 1 (factMarkOccXY) is a parameter for adjusting the extent of exclusion in *xy*. The *z* extent of the marked volume is large enough to contain the two SWC points: the distances from the top and bottom planes of the volume to the nearest SWC points are set to the maximum of *α_occ_r*_1_, *α_occ_r*_2_, or *z*_*occ*_ = 3 *µ*m (zOcc). Increasing the volume of the marked pixels prevents creations of spurious SWC points. However, if the volume is too large, correct SWC points can be eliminated, especially when branches are close to each other. These opposing constraints should guide the choice of the appropriate size for the volume.

The parameters for marking pixels occupied are in Table 6.

**Table 6:** Parameters for marking pixels near the SWC points occupied.

| Parameter | Name | Value | Meaning |
| --- | --- | --- | --- |
| $\alpha_{occ}$ | **factMarkOccXY** | 1 | factor for adjusting radius of the marked volume |
| $z_{occ}$ | **zOcc** | 3 $\mu\text{m}$ | lower bound for extend of the volume in $z$ |

### Thick dendrites and the soma

Thick dendrites and the soma can be missing from the mask created with the valley detectors. There are two main reasons: (1) their dimensions are much larger than the length scale of the valley detectors; (2) the intensities of the pixels inside them are uniform. Consequently, only the pixels at their boundaries show up in the mask. This leads to the error of no SWC points for these structures. To correct this problem, we check the existence of thick dendrite and the soma in each tiff stack. The check is based on the observation that these structures are typically well stained and the pixels in them are very dark.

Specifically, we create and compare two 2D projections of the tiff stack. In the first one, we only project the pixels that are marked occupied because they are in the vicinity of the existing SWC points. We enlarge the marked volume by increasing *α*_*occ*_ and *z*_*occ*_ to three times of the original values. This is to ensure that all pixels associated with the dendrites already covered the SWC points, including the shadows the dendrites in the out-of-focus planes, and completely marked. From this 2D projection we determine a threshold, which is set to the intensity of the darkest 5% of pixels. This threshold indicates the darkness of the pixels covered by the existing SWC points.

In the second 2D projection, we project only the pixels that are not marked occupied. We then count the number of pixels that are darker than the threshold determined from the first 2D projection. If this number is larger than the number of pixels in an area 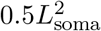, where *L*_soma_ = 2 *µm* (`somaLengthScale`) is the length scale of the soma, we decide that the tiff stack contains thick dendrite and/or the soma since there are significant number of dark pixels that are uncovered by the SWC points.

To place SWC points on thick dendrites and the soma, we create a 2D projection of the tiff stack with all pixels, smooth it, and create a mask by selecting top *θ_soma_* = 0.08 (somaSparseThr) fraction of the darkest pixels. From the mask we create SWC points as described before. SWC points are created only if their positions are not marked occupied by the existing SWC points, using the original values of *α_occ_* and *z*_*occ*_.

The parameters for creating SWC points for thick dendrites and soma are listed in Table 7.

**Table 7:** Parameters for creating SWC points for thick dendrites and the soma.

| Parameter | Name | Value | Meaning |
| --- | --- | --- | --- |
| $L_{soma}$ | **somaLengthScale** | 2 $\mu\text{m}$ | length scale of the soma |
| $\theta_{soma}$ | **somaSparseThr** | 0.05 | fraction for the threshold of creating mask |

### Extending SWC points in 3D

The SWC structures created from the masks are often incomplete, mostly due to the limitations of the masks in separating nearby branches in 2D projections. These branches could be well separated in the tiff stack although their 2D projections are not; therefore it is useful to extend the SWC structure using the tiff stack.

To minimize the interference from noise, we delete isolated SWC points that are not connected to any other SWC points. To ensure that the extension does not create duplicated SWC points, we mark pixels nearby the existing SWC points occupied (red circles, Fig. 7a). From an end SWC point (*x, y, z, r*) (yellow circle, Fig. 7a), we search the plane at *z* for the candidate for the next point. To reduce noisy fluctuations, the pixel intensity in the plane is smoothed with a Gaussian filter with *σ* = 2 pixels.

The search is done by finding a path starting from the end point and through the neurite (white line, Fig. 7a). To do so, we draw an arc (blue arc, Fig. 7a) of radius *r*_*search*_ = 3 *µ*m (`searchMax`), or (*α*_*s,m*_ + 2)*r*, whichever is greater. Here *α*_*s,m*_ = 1.2 (`factSearchMin`) is a factor. The arc is restricted to a range of angle (*θ*_thr_ = *π/*3, angle), where the angle is between the line from a pixel in the arc to the end point (black lines, Fig. 7a) and the line from the end point to its connected SWC point (yellow line, Fig. 7a). From each point on the arc, we compute the intensity weighted shortest distance path to the end point (black line, Fig. 7b). The resulting profile of the distance is Gaussian smoothed with parameter 2.0 (green line, Fig. 7b). The standard deviation of the difference between the original and the smoothed profiles is taken as the reference for the fluctuations in the distances. We then search for the significant local minima in the smoothed profile. To be significant, the local minimum must be smaller than a threshold set to the mean of the maximum and the minimum of the smoothed profile, minus *α*_*DD*_ times the standard deviation. Here *α*_*DD*_ = 20 (factSigmaDD) is a factor.

The shortest paths from the local minima to the end point are used to place the next SWC point. When there are multiple paths, each path is tested sequentially. The depth of the neurite along a chosen path is computed using the xy-path technique. Starting from the end point and following the path, we test placing a new SWC point *x*_*c*_, *y*_*c*_, *z*_*c*_, *r*_*c*_. We test the validity of the candidate SWC point, which also adjusts *x*_*c*_, *y*_*c*_ and *r*_*c*_. The validity test is done three times. If the distance from (*x*_*c*_, *y*_*c*_) to (*x, y*) is smaller than *α*_*s,m*_*r*, it is not accepted. If the candidate point passes all three tests, we check if it is near an existing SWC point; if so, we check whether the existing SWC point is connectable to the end point. If connectable, the existing SWC point is added to the candidate pool for connecting the end point to the existing SWC points. If a valid candidate point does not come close to the existing SWC points, and it is connectable to the end point, we create a new SWC point at (*x_c_, y_c_, z_c_*) with radius *r*_*c*_. It is connected to the end point, and serves as a new end point from which the extension continues. If no new SWC point is created after checking all paths, we check the list of candidates for connections, and select the one with the minimum distance and connect it to the end point.

The parameters used in extending SWC points are listed in Table 8.

**Table 8:**
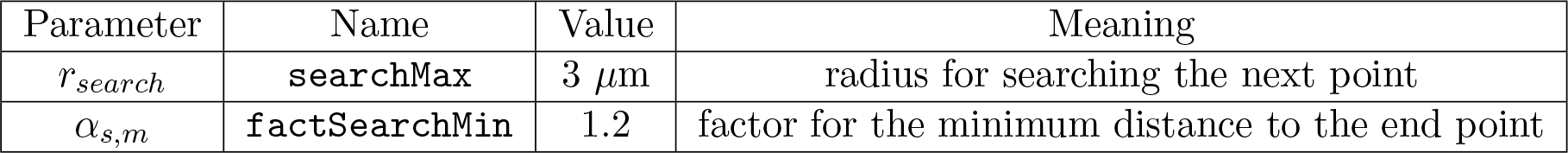
Parameters for extending SWC points in 3D.

### Connecting broken segments

After extending SWC points in 3D, a continuous branch can still be represented with broken segments of SWC points, especially if the underlying signal is weak or there are closely crossing branches (Fig. 2b,c). We connect these segments with heuristic rules to recover the branch continuity. To do so, we compute the Euclidean distance between all pairs of end points that are not connected and the differences in *z* are within the allowed range as described before. If the distance of the pair in *xy* is smaller than 1.5(*r*_1_ + *r*_2_), where *r*_1_, *r*_2_ are the radii, they are judged to be close to each other and are connected.

After connecting the nearby pairs, we consider more distant ones. If the two end points are within *α_xy_*(*r*_1_ + *r*_2_) in *xy*, where *α_xy_* = 2 (distFactConn) is a factor; and if the angles between the two lines, linking the end points to their respective connected SWC points, is smaller than *θ_thr_* = *π/*3 (angle); then the two points are preserved in the candidate pool for potential connections. We then iteratively connect the pairs of end points in the pool, connecting the closest available pairs first. Once connected, the end points are excluded from further connections. This pairwise connection stops if the pairs are all considered or if further connections create loops in the SWC structure.

The parameters used in connecting end points are listed in Table 9.

**Table 9:** Parameters for connecting end points.

| Parameter | Name | Value | Meaning |
| --- | --- | --- | --- |
| $\alpha_{xy}$ | **distFactConn** | 2 | factor for judging the closeness of end points |

### Subdividing tiff stack in *z*

The 2D projection can be complicated when there are many branches in one stack. This often leads to occlusions in the 2D projection and missed branches in the reconstruction. One way of mitigating this problem is to divide the tiff stack in *z* into *n*_*div*_ = 8 (nSplit) slabs with equal heights in *z*. We create SWC points separately for each slab. When creating the 2D projection for a slab, we include extra volume in *z* by extending the height in both directions by *z*_*ext*_ = 3 *µ*m (zext). This is useful for getting good projections of branches that are near the dividing planes between the slabs. The *z*-image also includes the extended volume. Any SWC points whose depths are beyond the slab boundary are deleted. The extension from the end points are done with the entire tiff stack, which helps to connect SWC points that belong to the same branch but are cut by the subdivision.

The parameters of subdivision are listed in Table 10.

**Table 10:** Parameters for subdividing a tiff stack.

| Parameter | Name | Value | Meaning |
| --- | --- | --- | --- |
| $n_{div}$ | **nSplit** | 8 | number of subdivisions |
| $z_{ext}$ | **zext** | 3 $\mu\text{m}$ | extension of sub-slabs when creating SWC points |

### Combining SWCs for the entire neuron

The image of an entire neuron consists of multiple tiff stacks stitched together (Fig. 10a). The coordinates of the stacks relative to the first stack are determined during the stitching. For each stack we obtain the SWC structure, and shift the positions of the SWC points by the relative coordinates of the stack. The SWC points of individual stacks are read-in sequentially. To avoid duplicated SWC points in the overlapping regions of adjacent stacks, pixels near the SWC points that are already created are marked occupied by setting the parameters *z*_*occ*_ 5 times of the usual value and *r*_*occ*_ 2 times. If the position of SWC points are at the marked pixels, they are deleted. For individual stacks, the step of connecting the end points is omitted. Instead, after reading in the SWC points of all stacks, we extend the SWC points from the end points, and then connect the new end points. To eliminate noise, we delete very short leaf branches in the SWC structure (those that have fewer than *n*_*dmin*_ = 5 SWC points, `minNumPointsBr`). In addition, we eliminate isolated branches that are shorter than *l*_*min,iso*_ = 20 *µ*m (`minLenBrIso`). Increasing this number reduces noise in the reconstruction, but can also delete some of the correct reconstruction. The reconstructed SWC structure for the example neuron is shown in Fig. 10b-d.

The parameters are listed in Table 11.

**Table 11:** Parameters for reducing noise in SWC structure.

| Parameter | Name | Value | Meaning |
| --- | --- | --- | --- |
| $n_{dmin}$ | **minNumPointsBr** | 5 | minimum number of SWC points allowed in leafs |
| $l_{min,iso}$ | **minLenBrIso** | 20 $\mu\text{m}$ | minimum length allowed in isolated branches |

